# Increasing plasma bile salt levels with Bulevirtide alleviates DSS-induced colitis and LPS-induced inflammation

**DOI:** 10.64898/2026.04.21.719641

**Authors:** Thuc-Anh Nguyen, Reinout LP Roscam Abbing, Pim J Koelink, Joost M Lambooij, Wietse In het Panhuis, Dirk R de Waart, Isabelle Bolt, Suzanne Duijst, Esther Vogels, Ricky Siebeler, Menno PJ de Winther, Bruno Guigas, Manon E Wildenberg, Coen C Paulusma, Stan FJ van de Graaf

## Abstract

**Background & Aims:** Bulevirtide, a viral entry inhibitor used to treat chronic hepatitis delta virus (HDV) infection, targets the hepatic bile salt transporter Na^+^-Taurocholate Co-transporting Polypeptide (NTCP). As Bulevirtide displays preclinical potential to mitigate cholestatic liver injury, NTCP inhibition is currently explored as treatment for primary sclerosing cholangitis (PSC), a condition frequently associated with colitis. Here, we investigated the immunomodulatory effects of Bulevirtide in lipopolysaccharide (LPS)-induced inflammation and dextran sodium sulfate (DSS)-induced colitis in mice.

**Methods:** The immunomodulatory properties of the bile salt taurochenodeoxycholic acid (TCDC) were investigated in LPS-challenged mouse bone marrow-derived macrophages (BMDM) and human BLaER1 macrophages. The therapeutic efficacy of Bulevirtide against LPS-induced inflammation and DSS-induced colitis was evaluated in *Slco1a/1b^-/-^* FVB and C57BL/6J mice, which recapitulate human bile salt dynamics.

**Results:** In BMDMs, TCDC reduced pro-inflammatory tumor necrosis factor alpha (TNF), increased anti-inflammatory interleukin (IL)-10, and suppressed inflammasome activation, as evidenced by reduced IL-1β, IL-18 and cleaved-IL-1β levels. Consistently, TCDC also reduced *TNF* and *IL1B* expression in human BLaER1 macrophages. In both FVB and C57BL/6J *Slco1a/1b^-/-^ mice*, Bulevirtide increased plasma bile salt levels at least 30-fold. This systemic elevation of bile salts reduced plasma TNF and increased IL-10 in LPS-treated mice. Moreover, Bulevirtide attenuated DSS-induced colitis, evidenced by reduced disease scores and reduced intestinal *Tnf* expression.

**Conclusion:** These findings highlight the anti-inflammatory effects of bile salts in preclinical models of colitis and support NTCP inhibition as a future therapeutic strategy to ameliorate both cholestasis and colitis in PSC.

**Graphical abstract:** 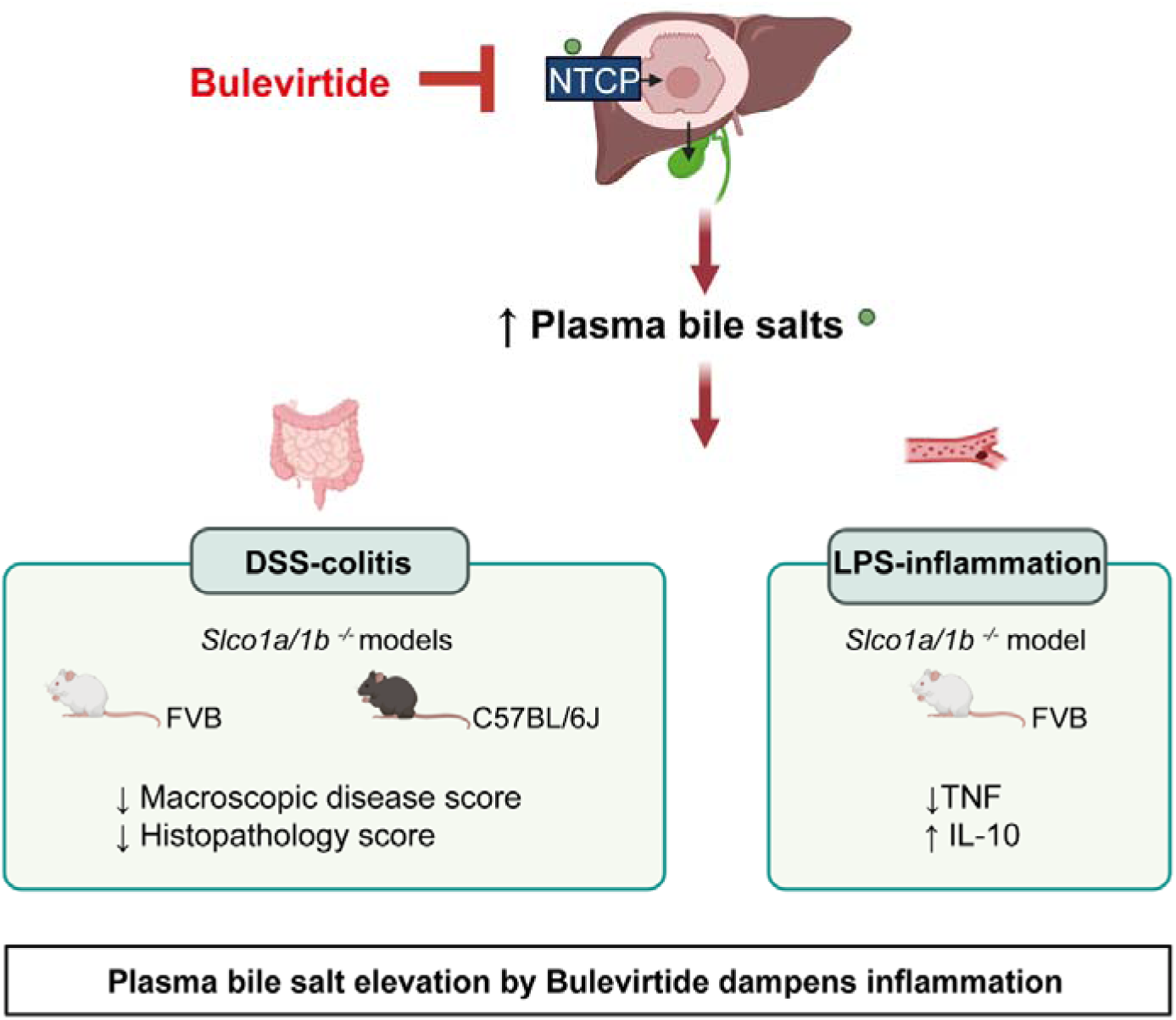

**Synopsis:** Inhibition of the Na^+^-Taurocholate Co-transporting Polypeptide using Bulevirtide induces systemic bile salt elevation and mitigates acute inflammation and colitis in mice. These findings support clinical evaluation of Bulevirtide in primary sclerosing cholangitis with protective effects against cholestasis and colitis.

## Introduction

Bile salts not only facilitate the digestion and absorption of dietary lipids and fat-soluble vitamins, but also act as signaling molecules that regulate metabolism and inflammation (1). Bile salt treatment decreases production of the pro-inflammatory cytokines tumor necrosis factor alpha (TNF) and interleukin (IL)-1β in macrophages (2–4). These effects may be mediated by activation of the bile salt responsive receptors Takeda G-protein-coupled Receptor 5 (TGR5) and nuclear receptor Farnesoid X receptor (FXR) (5, 6). In cholestatic liver disease, bile salt accumulation in the liver causes hepatocellular damage and inflammation, which is mainly attributed to toxic hydrophobic bile salt species (7). In contrast, the hydrophilic ursodeoxycholic acid (UDCA) mitigates cholestatic liver injury and serves as the first-line therapy for primary biliary cholangitis (PBC) (7).

Primary sclerosing cholangitis (PSC) is a rare cholestatic disease characterized by chronic inflammation and fibrosis of the intra- and extrahepatic bile ducts (8). There is currently no approved treatment for PSC. Up to 80% of PSC patients also suffer from ulcerative colitis (UC) (9). Patients with PSC-UC exhibit a distinct phenotype compared to classical UC, with inflammation predominantly in the cecum and ascending colon, showing milder symptoms, but a significantly higher risk of developing colorectal cancer and cholangiocarcinoma than patients with PSC or UC alone (9). Therefore, it is necessary to develop safe and effective therapies for PSC targeting both cholestatic liver injury and associated UC.

Previously, we demonstrated that inhibition of the Na^+^-Taurocholate Co-transporting Polypeptide (NTCP) by Bulevirtide (previously Myrcludex B) attenuated cholestatic liver injury in mice, by i) reducing hepatocellular bile salt accumulation, while concomitantly increasing systemic bile salt levels and ii) increasing the phospholipid to bile salt ratio in bile, rendering less toxic bile (10). NTCP is the main hepatic uptake transporter for conjugated bile salts that re-enter the liver via the portal blood. In healthy individuals, Bulevirtide administration transiently increases plasma bile salt levels to around 200 µM (11–13). This elevation is well tolerated, and NTCP inhibition by Bulevirtide has proven safe for treating individuals with chronic infection with HBV and HDV, of which NTCP is the viral entry receptor (12, 14). In contrast to humans, mice do not demonstrate systemic bile salt elevation upon NTCP inhibition, due to relative high hepatic bile salt uptake by the Organic Anion Transporting Polypeptide (OATP) 1a/1b members (15). In *Slco1a/1b^-/-^ mice*, Bulevirtide induced systemic plasma bile salt elevation that resembled the human bile salt dynamics (16). Importantly, 48-week Bulevirtide therapy in patients with HBV/HDV coinfection not only reduced the number of infected hepatocytes but also suppressed the expression of inflammatory genes in liver biopsies (17). These findings suggest that Bulevirtide exerts immunosuppressive effects. Whether this is entirely due to the reduction of viral titers or include a direct immunomodulatory effect of increased plasma bile salt levels remains unclear.

Here, we explored the immunomodulatory properties of bile salts and investigated whether Bulevirtide-induced plasma bile salt elevation could alleviate colitis. Our data demonstrate that bile salts suppressed inflammation in LPS-stimulated bone marrow-derived macrophages (BMDMs) and human macrophages. Bulevirtide-mediated elevation of plasma bile salt levels diminished LPS-induced inflammation in *Slco1a/1b^-/-^ mice*. Furthermore, Bulevirtide alleviated dextran sodium sulfate (DSS)-induced colitis, as shown in two different *Slco1a/1b^-/-^* mouse models.

## Results

### TCDC suppresses inflammation in murine BMDMs and human macrophages

Cytokine-driven inflammation is central to colitis, with macrophages serving as a major source of these mediators (18). Therefore, we evaluated the immunomodulatory effects of bile salts on macrophages *in vitro*. BMDMs were treated with TCDC at 100 µM and activated with LPS; ATP was given to activate cytokine secretion (Fig. 1A). TCDC administration reduced gene expression of *Tnf* (−48%, p=0.0286) and *Il6* (−39%, p=0.0286), whereas *Il1b, Mcp1*, *Il12a* and *Il10* expression tended to be reduced (Fig. 1B). In line with reduced mRNA expression, TNF in the medium of LPS-stimulated BMDMs was also reduced by TCDC (−35%, p=0.0286) (Fig. 1C). Interestingly, secretion of the anti-inflammatory cytokine IL-10 tended to be increased by TCDC (+61%, p=0.1000) (Fig. 1D). Next, we examined whether TCDC affected inflammasome activation. Protein analysis of BMDM whole cell lysates indicated that TCDC treatment did not affect intracellular pro-IL-1β levels but clearly reduced cleaved-IL-1β levels compared to non-treated control (Fig. 1E). This suggested reduced caspase-1 activity in TCDC-incubated BMDMs, which was confirmed by diminished cleavage of pro-caspase-1 to its active p20 form (Fig. 1E). Correspondingly, TCDC tended to reduce the secretion of the inflammasome dependent cytokines IL-1β (−36%, p=0.0649) (Fig. 1F) and IL18 (−88%, p=0.0022) (Fig. 1G). Overall, these data indicate that TCDC at 100 µM dampened the LPS-induced inflammatory response in BMDMs.

**Fig. 1.**
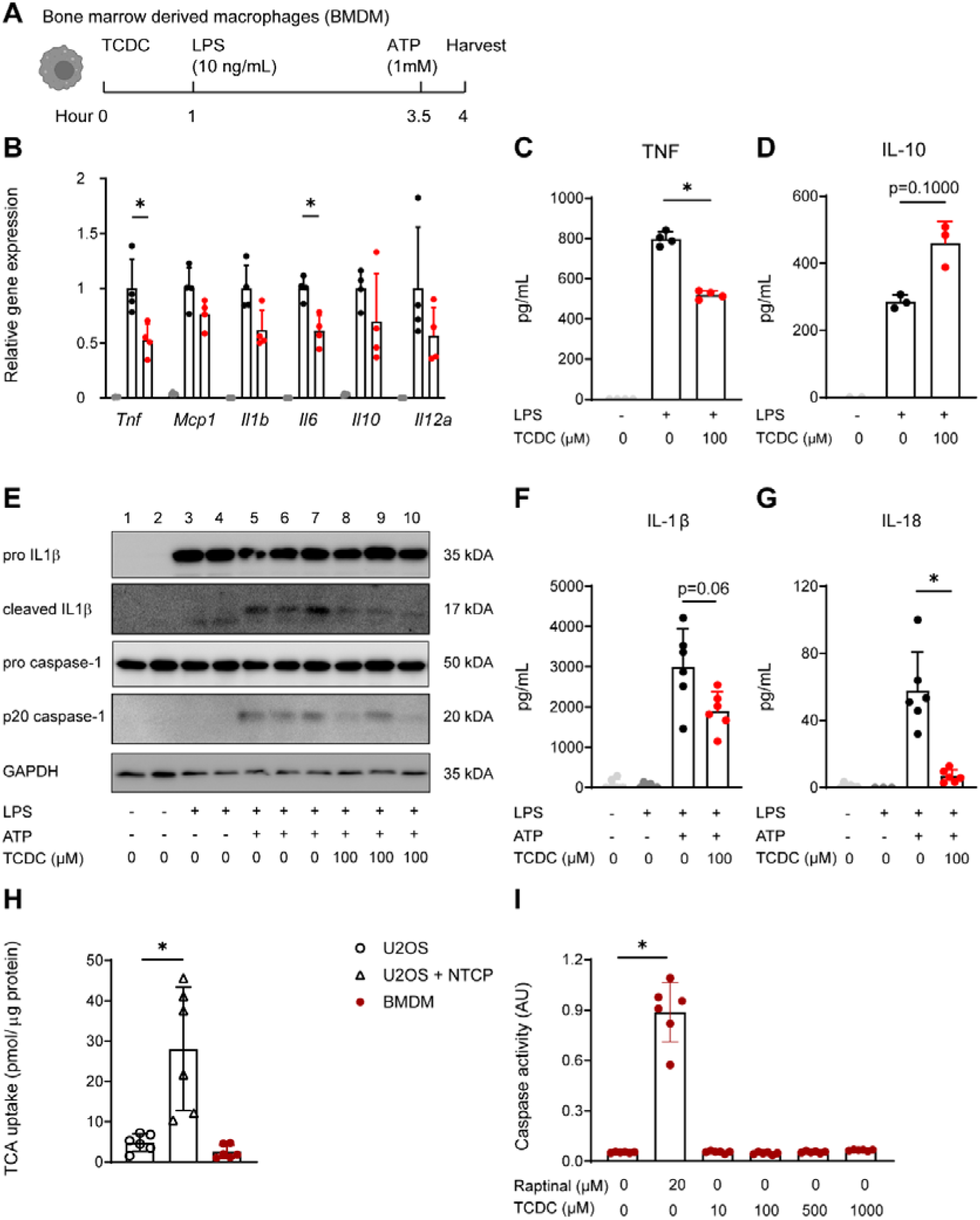
TCDC has immunosuppressive effects on BMDMs. (A) Experimental schematic; BMDMs were treated with TCDC (100 µM) for 1 hour, followed by LPS (10ng/mL) for 3 hours. To activate the inflammasome, cells were also treated with ATP (1mM, 30 minutes before harvesting). (B) Gene expression of the cytokines TNF, MCP1, IL-1β, IL-10 and IL-12 in BMDMs. (C) TNF and (D) IL10 concentration in medium of BMDMs was measured by ELISA. (E) Protein levels of IL-1β and caspase-1 were measured in whole cell lysates of BMDMs. (F) IL-1β and (G) IL-18 concentrations in medium of BMDMs were measured by ELISA. (H) TCA uptake of BMDMs, in comparison to U2OS parental and U2OS-NTCP cells. (I) Caspase 3/7 activity was assessed as a marker of apoptotic activity in BMDMs incubated with the indicated concentrations of TCDC. n=3-6 per group. Experiments were performed 3 times and representative data are shown. Data are means ± SD. Significance was assessed with Mann-Whitney U test. *\* p*<0.05.

To exclude that the observed effects were caused by TCDC-induced toxicity, we investigated bile salt uptake and TCDC-induced apoptosis in BMDMs. BMDMs did not show any uptake of the bile salt taurocholic acid (TCA) (Fig. 1H), as levels were comparable to U2OS cells, which lack bile salt uptake transporters (19). U2OS cells expressing NTCP served as positive control and clearly displayed TCA uptake. TCDC treatment for 3 hours did not induce caspase 3/7 activity in BMDMs, even at high concentrations of 500 µM or 1000 µM (Fig. 1I). Therefore, the TCDC-induced reduction in inflammatory responses in BMDMs was not caused by intracellular accumulation of bile salts and TCDC-induced toxicity.

To examine whether the immunosuppressive effects of TCDC were conserved between species, we studied the effects of TCDC on LPS-induced inflammation in human macrophage BlaER1 cells. Cells were treated with several concentrations of TCDC, followed by LPS and ATP stimulation (Fig. 2A). Compared to control, TNF gene expression tended to reduce with TCDC at 100 µM (−20%, p= 0.0547) and was significantly reduced at 200 µM (−32%, p=0.0169) (Fig. 2B). IL1B gene expression was also decreased by TCDC, most strongly at 200 µM (−27%, p=0.0157) (Fig. 2C). No effects on IL10 gene expression were observed (Fig. 2D). Similar to murine BMDMs, we found that cell viability of BlaER1 cells was not impaired by the TCDC treatment (Fig. 2E). These results demonstrate that TCDC induces immunosuppressive effects in mouse and human macrophages.

**Fig. 2.**
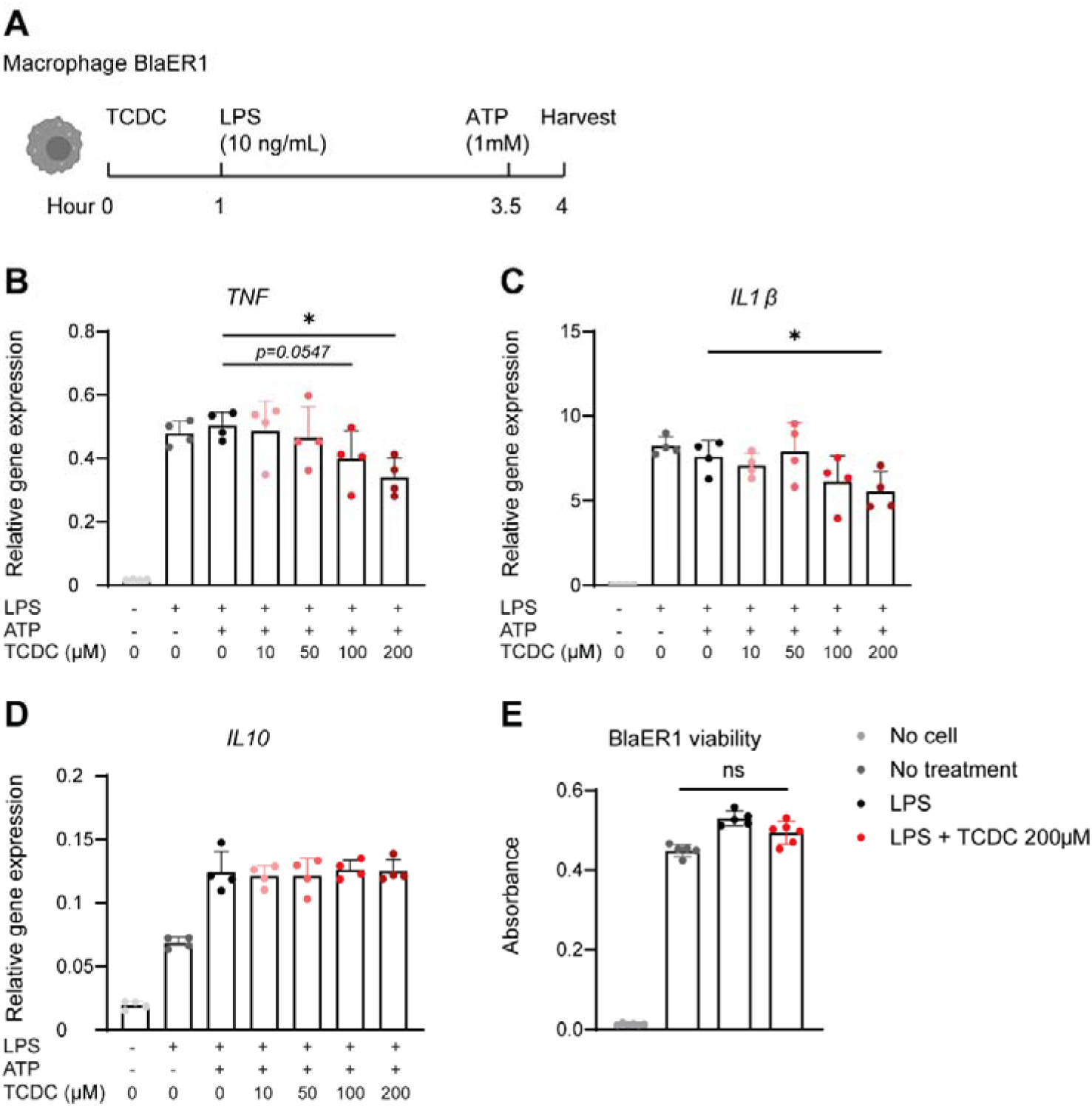
TCDC has immunosuppressive effects on human macrophages. (A) *In vitro* experimental schematic; BlaER1 macrophages were treated with TCDC, followed by LPS (10ng/mL) and ATP (1mM) stimulation. Gene expression of cytokine TNF (B), IL-1β (C) and IL-10 (D) in BlaER1 cells. (E) Viability of BlaER1 cells treated with TCDC for 1 hour, followed by LPS stimulation. Data are means ± SD. Statistical significance was assessed with Kruskal-Wallis test in (B, C, D, E). * *p<*0.05; ns, not significant (pL>L0.05).

### Systemic bile salt elevation by Bulevirtide attenuates LPS-induced inflammation in mice

LPS triggers acute systemic inflammation, marked by elevated levels of cytokines (20). We explored whether Bulevirtide-induced elevation of systemic bile salt levels could alleviate acute systemic inflammation in *Slco1a/1b* cluster knockout FVB mice (21), hereafter called *Slco1a/1b^-/-^* FVB mice (Fig. 3A). NTCP inhibition led to a substantial increase in plasma bile salt levels by ∼25-fold (p=0.0007) (Fig. 3B) and a shift in the bile salt pool composition from unconjugated to taurine-conjugated bile salts (Fig. 3C). In particular, TCA and tauro-β-muricholic acid (TβMC) levels were elevated (Fig. 3C). Notably, the high systemic bile salt concentrations coincided with reduced plasma TNF (−36%, p=0.0401) (Fig. 3D) and increased plasma IL-10 levels (+140%, p=0.0012) (Fig. 3E). Plasma levels of IL-1β, IL-6, IL-12 and MCP1 were not affected (Fig. 3F-I). These results suggest that elevation of systemic bile salt levels *in vivo* alleviated an inflammatory response upon an acute LPS-induced insult.

**Fig. 3.**
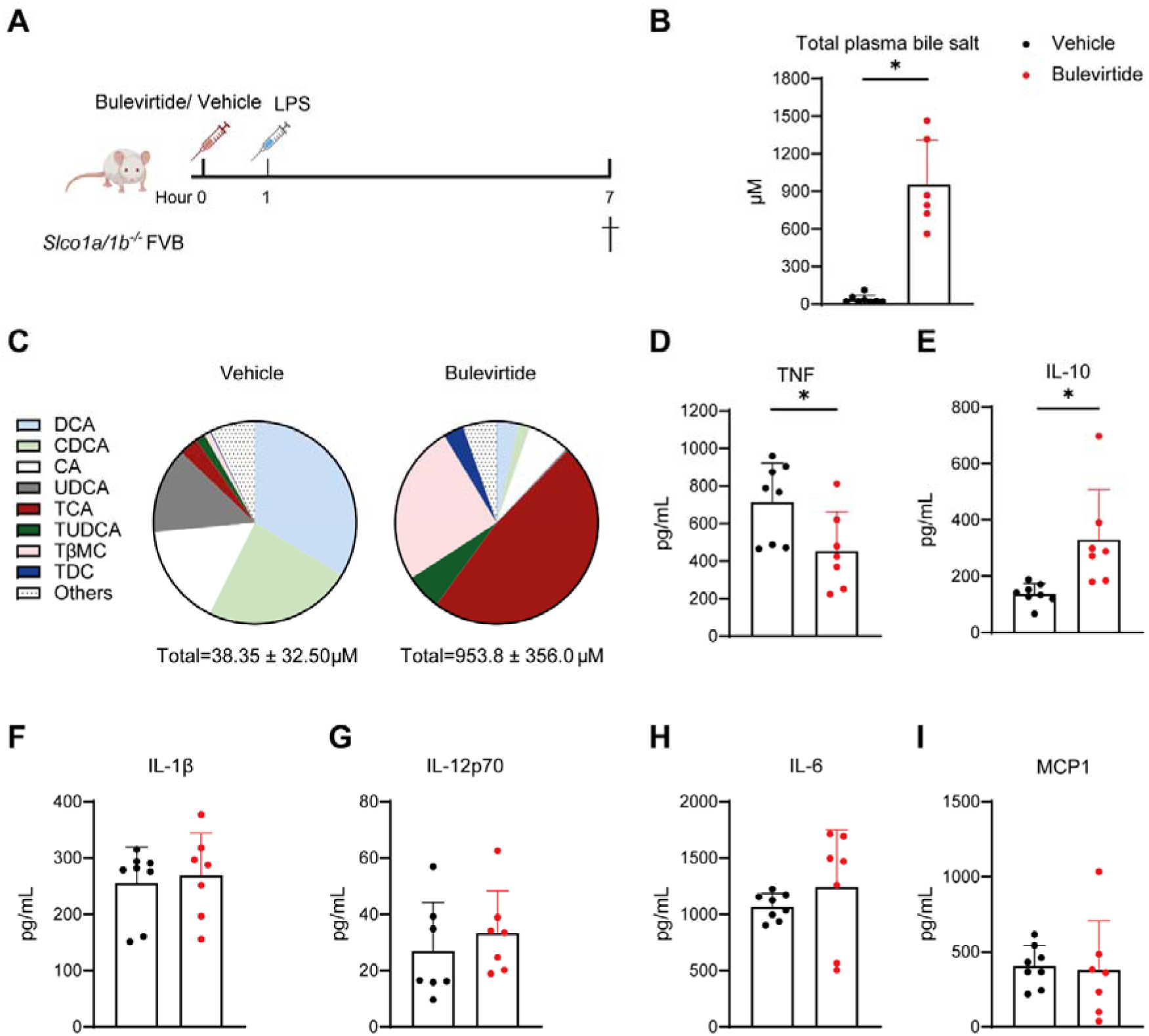
Systemic bile salt elevation by Bulevirtide attenuates inflammatory responses to LPS *in vivo*. (A) Schematic experimental overview; *Slco1a/1b^-/-^* FVB mice were injected with vehicle or Bulevirtide 2.5 µg/g, followed by LPS injection at 20 µg/g. (B) Total bile salt concentration and (C) concentration of individual bile salt species measured by HPLC. Plasma concentration of the cytokines TNF (D), IL-10 (E), IL-1β (F), IL-12p70 (G), IL-6 (H) and MCP1 (I). IL-1β was measured by ELISA, whereas other cytokines were measured by cytometric bead array. n=8 per group. Data are means ± SD. Significance was assessed with Mann-Whitney U test. *\* p*<0.05.

### Systemic bile salt elevation by Bulevirtide attenuates colitis in FVB mice

Next, we examined whether Bulevirtide-induced elevation of systemic bile salt levels could reduce inflammation in DSS-induced colitis (Fig. 4A). Since FVB mice appeared somewhat resistant to DSS (Fig. S2), we used 6% DSS to induce colitis. WT mice on normal drinking water had low plasma bile salt levels, mainly consisting of deoxycholic acid (DCA), TCA and UDCA, which increased following DSS exposure (∼2-fold, p=0.0249) (Fig. 4B, Fig S3A). Compared to the DSS-treated WT mice, DSS-treated *Slco1a/1b^-/-^* FVB mice with concomitant daily injections with vehicle demonstrated further elevation of baseline plasma bile salt levels (∼6-fold, p=0.0035). Bulevirtide resulted in an additional 23-fold increase in plasma bile salts relative to vehicle in DSS-treated *Slco1a/1b^-/-^* FVB mice (p=0.0035) (Fig. 4B). Similar to the acute LPS model (Fig. 3C), Bulevirtide induced a shift toward a taurine-conjugated bile salt pool composition, driven by significantly increased concentrations of TβMC, TCA and cholic acid (CA) (Fig. 4C). DSS/vehicle-treated WT and *Slco1a/1b^-/-^*FVB mice had reduced colon length and increased colon density, indicating colonic inflammation (Fig. 4D). Macroscopic disease scores were elevated in the DSS-treated mice (Fig. 4E), as were the histopathology scores of the colon, evidenced by thickening of the sub-mucosa, massive immune cell infiltration, and epithelial membrane damage (Fig. 4F-G). Notably, Bulevirtide treatment markedly ameliorated colonic inflammation, evidenced by restored colon length (Fig. 4D) as well as significantly reduced macroscopic and histological disease scores compared to the vehicle group (Fig. 4E-F). Of note, DSS water intake was reduced in the Bulevirtide group (Fig. S3B). Despite this, DSS caused (moderate) body weight-loss (Fig. 4H) as well as an increased spleen weight in all groups (Fig. S3C), indicating that the DSS treatment promoted systemic inflammation in all groups.

**Fig. 4.**
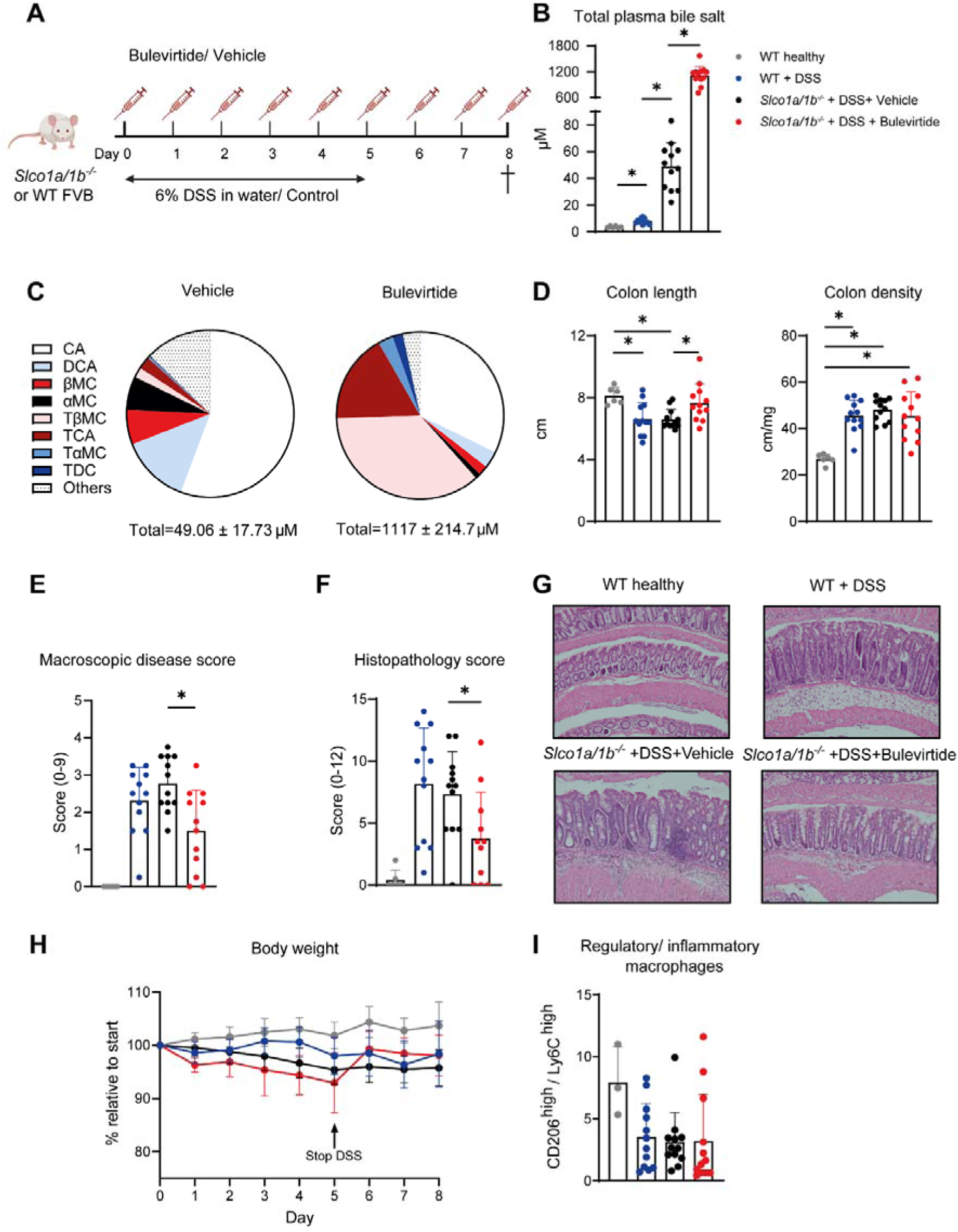
Systemic bile salt elevation by Bulevirtide attenuates DSS-induced colitis in FVB mice. (A) Schematic experimental overview; Colitis was induced in WT and *Slco1a/1b^-/-^*FVB mice by 6% DSS in drinking water for 5 days. Mice were treated with vehicle or Bulevirtide 2.5 µg/g daily. (B) Total bile salt concentration and (C) Concentration of individual bile salt species measured by HPLC. (D) Colon length (cm) and density (mg/cm). (E) Macroscopic disease score at sacrifice. (F, G) Histopathology score of colon at the end of treatment. (H) Body weight relative to start of DSS. (I) Ratio between regulatory and inflammatory subsets of macrophages in colon measured by flow cytometry. Data are means ± SD. Statistical significance was assessed with Kruskal-Wallis test in (B, D, E, F, I). * *p<*0.05.

To assess whether high plasma bile salt levels affected colonic macrophage polarization and thereby could contribute to disease improvement, we performed flow cytometry analysis of colon-derived macrophages. DAPI^−^CD45^+^CD11b^+^Ly6G^−^ myeloid cells were evaluated for the expression of Ly6C and CD206, markers for inflammatory and regulatory subsets, respectively (18). Bulevirtide treatment did not increase the ratio of regulatory CD206^hi^ Ly6C^lo^ macrophages over inflammatory CD206^lo^ Ly6C^hi^ macrophages in the colon (Fig. 4I), suggesting that bile salts interfere at a different level in FVB mice.

### Systemic bile salt elevation by Bulevirtide attenuates colitis in C57BL/6J mice

As a second model to investigate the immunomodulatory effects of plasma bile salt elevation by Bulevirtide in DSS-induced colitis, we used mice with a C57BL/6J genetic background, which are more susceptible to DSS-induced colitis than FVB mice (22). To generate a C57BL/6J mouse model that recapitulates human bile salt uptake dynamics as observed in *Slco1a/1b^-/-^* FVB mice, we inactivated *Slco1a1, Slco1a4 and Slco1b2* in C57BL/6J mice (hereafter called *Slco1a/1b^−/−^* B6) using CRISPR-Cas9 (described in Material and Methods) (Fig. S1A). Immunoblotting confirmed the loss of OATP1A1, 1A4 and 1B2, while NTCP appeared to be slightly upregulated (Fig. S1C). Mice exhibited normal growth and behavior. Furthermore, we examined plasma and biliary bile salt kinetics following NTCP inhibition with Bulevirtide in *Slco1a/1b^−/−^* B6 mice. All vehicle-treated WT and knockout strains showed rapid plasma clearance of [^3^H]TCA, which was paralleled by fast biliary excretion (Fig. S1D). Bulevirtide did not significantly affect the clearance rates in WT and *Slco1a1^-/-^* B6 mice, however, strongly delayed plasma clearance and biliary secretion of [^3^H]TCA in *Slco1a/1b^−/−^* B6 mice (Fig. S1D), indicating that hepatic bile salt uptake was effectively inhibited in this model.

Drinking water with 2.5% DSS was given to *Slco1a/1b^−/−^* B6 mice for 5 days, together with daily injections of Bulevirtide or vehicle (Fig. 5A). A control group that received water only was also included. Both vehicle and Bulevirtide groups exhibited a similar trend in water intake (Fig. S4A). NTCP inhibition markedly increased plasma bile salt levels by more than 30-fold compared to vehicle (p<0.0001) (Fig. 5B), and (similar to the previous models) changed the bile salt pool composition (Fig. 5C). This elevation was mainly due to increased levels of CA and taurine-conjugated species, including TCA and TβMC. Both groups showed gradual body weight loss upon DSS challenge (Fig. 5D). Bulevirtide attenuated DSS-induced body weight loss (Fig. 5D) and prevented DSS-induced shortening of colon length (+9%, p = 0.0382), while colon density remained unchanged (Fig. 5E). The macroscopic disease score was decreased in the Bulevirtide group (−24%, p=0.0221), indicating lower intestinal inflammation / edema (Fig. 5F). Similarly, the histopathology score was reduced in the Bulevirtide group (−12%, p=0.0347) (Fig. 5G), as indicated by less immune cell infiltration and epithelial damage (Fig. 5H). Bulevirtide reduced intestinal gene expression of *Tnf* (−13%, p=0.0263), however, did not affect *Il1b* and *Il10* expression (Fig. 5I). These results show that elevation of bile salt levels by Bulevirtide alleviated DSS-induced colitis in C57BL/6J mice, in line with FVB mice.

**Fig. 5.**
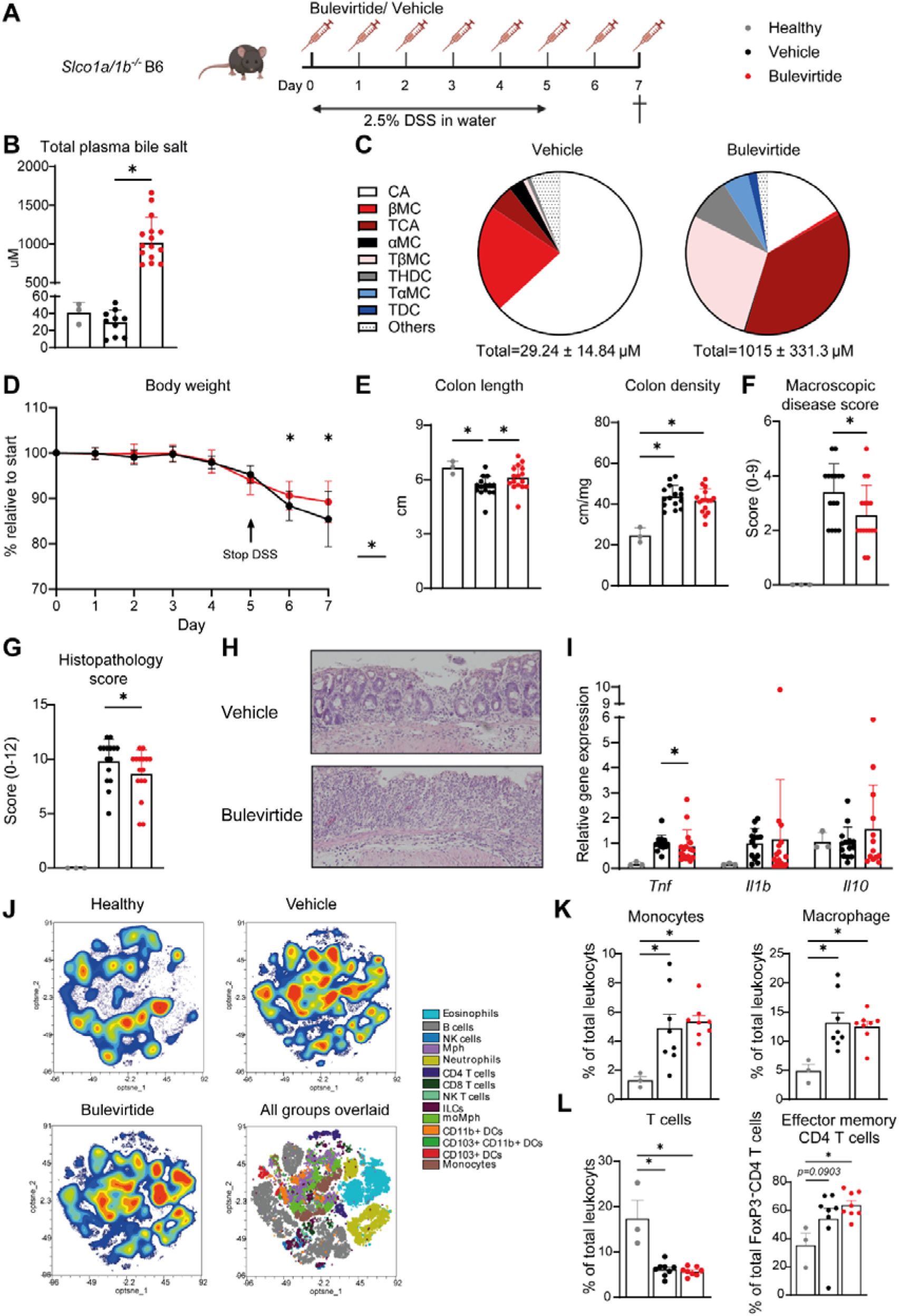
Systemic bile salt elevation by Bulevirtide attenuates DSS-induced colitis in C57BL/6J mice. (A) Schematic experimental overview; colitis was induced in *Slco1a/1b^-/-^* B6 mice by 2.5% DSS in drinking water for 5 days. Mice on normal drinking water were included as healthy controls. DSS-mice were treated with vehicle or Bulevirtide 2.5 µg/g daily. (B) Total bile salt concentration and (C) Concentration of individual bile salt species measured by HPLC. (D) Body weight relative to start of DSS. (E) Colon length and density, and (F) macroscopic disease score at sacrifice. (G, H) Histopathology score of colons at the end of treatment. (I) Colonic gene expression of cytokines TNF, IL-1β and IL-10. (J) Optimized t-Distributed Stochastic Neighbor Embedding (optsne) used to visualize global changes in immune composition of the colon in healthy, vehicle and Bulevirtide treated samples with each plot showing pooled data from all mice per group. (K) Monocytes, macrophages and (L) Total T cells presented as frequency of the total leukocyte pool in addition to effector memory CD4 T cells presented as frequency of total FoxP3^-^ CD4 T cells. Data are means ± SD. Statistical significance was assessed with Kruskal-Wallis test in (B, E, F, G, I, K, L), Mann-Whitney test in (D). * *p<*0.05.

To investigate which immune cell types could be important in the Bulevirtide-mediated alleviation of intestinal inflammation, we performed flow cytometry analyses on leukocytes isolated from colon, mesenteric lymph nodes (mLNs), and spleen. In colon, DSS supplementation resulted in infiltration of immune cells (Fig. 5J), particularly monocytes, macrophages and neutrophils (Fig. 5K, Fig. S6A). The total T-cell population was reduced (Fig. 5L), with an increase in the CD4^+^ T-cell proportion and no change in the CD8^+^ T-cell proportion (Fig. S6C). Further analyses demonstrated that the effector memory CD4^+^ T-cell proportion tended to increase (Fig. 5L). Also, in mLNs, the proportions of monocytes, macrophages, eosinophils (Fig. S7B), central memory CD4^+^ T-cells (Fig. S7D) were increased. The increase in effector and innate immune cell compositions in both the colon and mLNs indicated inflammation induced by DSS. Furthermore, RORγt^+^ CD4^+^ T-cells, which are typically increased in response to DSS (23, 24), were also increased in the mLNs (Fig. S7D), but tended to decrease in the colon (Fig. S6C). DSS supplementation also resulted in a reducing trend in colonic RORγt^+^Helios^-^Tregs (regulatory T cells) (Fig. S6C). There was, however, no effect of Bulevirtide on inflammatory cell compositions compared to vehicle (Fig. 5J-L, Fig.S6-7). Additionally, general immune populations in the spleen were not affected by either DSS or the treatments, showing that inflammation in DSS colitis model is localized to the intestine (Fig. S8). Although spleen weight increased with DSS, it was also not affected by the treatments (Fig. S4B). These data suggest that Bulevirtide alleviated colitis and suppressed inflammatory cytokine expression without affecting immune cell compositions infiltrating the colon and mLNs.

## Discussion

Using mouse models, we investigated Bulevirtide – a safe and approved drug for chronically HDV infected individuals – as a potential pharmacological therapy for PSC-colitis and showed anti-inflammatory effects of elevated levels of systemic bile salts. Conjugated bile salts lowered pro-inflammatory TNF levels, increased the regulatory cytokine IL-10 levels, and suppressed inflammasome activation, in LPS-activated BMDMs. Similarly, bile salts dampened inflammation in BlaER1 human macrophages at concentrations reached in Bulevirtide-treated individuals. Similar results were seen *in vivo*, as systemic plasma bile salt elevation by Bulevirtide suppressed LPS-induced inflammation in FVB mice. Furthermore, Bulevirtide alleviated DSS colitis severity in two distinct mouse models.

These results reveal a potential therapeutic effect of Bulevirtide in colitis, in addition to its established anti-viral function and preclinically demonstrated efficacy to reduce cholestatic liver damage (10). In patients with HBV/HDV coinfection, Bulevirtide effectively improved liver damage and reduced HDV infection markers, with the greatest reduction observed at higher doses and longer treatment durations (12, 17, 25). Liver biopsies at baseline and after 48-week Bulevirtide treatment in humans displayed downregulation of pro-inflammatory genes including *TNF* and *IL1B*, as well as anti-viral *IFNG*, while still reducing the number of HDV-infected cells. This indicates that the anti-inflammatory effects of Bulevirtide did not interfere with its anti-viral function of blocking HDV entry. Furthermore, our previous study found that Bulevirtide dampened cholestatic liver injury *in vivo* by reducing hepatocellular bile salt accumulation and lowering bile toxicity (10). Also, pro-inflammatory cytokine- and pro-fibrotic genes were downregulated (10). Similarly, inhibiting the apical sodium-dependent bile salt transporter (ASBT), which mediates bile salt uptake from the intestinal lumen, reduced hepatic bile salt load and alleviated cholestasis in mice (26). However, in contrast to NTCP inhibition, ASBT inhibitors are associated with diarrhea, which is a significant concern in patients with colitis. (10, 26).

Importantly, Bulevirtide demonstrates an excellent safety profile in clinical settings. As plasma bile salt levels increase in patients with cholestatic liver diseases, a causal role as pruritogens has been postulated, although this appears to be incorrect (27). Bulevirtide treatment for 96 weeks to 3 years was effective against HBV/HDV co-infection, with no apparent correlation between pruritus and plasma bile salt elevation (12, 25). NTCP deficiency is relatively common in South-East Asia (28). Despite their high plasma bile salt levels, even at concentration up to 1500 µM, these individuals are asymptomatic, including lack of pruritus (10, 25). In the presence of physiological levels of albumin, bile salts do not activate known itch receptors (29). This suggests that elevated bile salt levels in plasma, induced by Bulevirtide, are unlikely to cause pruritus in PSC patients.

NTCP inhibition not only elevates bile salt levels, but also alters bile salt composition. In WT mice, bile salt concentrations in plasma are usually low and consist of mainly DCA, TCA, UDCA and some MC species in both FVB (Fig. S3A) and C57BL/6J strains (30). Ablation of *Slco1a/1b* increased basal plasma bile salt levels, with the pool dominated by unconjugated bile salts in both strains (Fig. 4C, Fig 5C) (15). NTCP inhibition by Bulevirtide leads to elevated levels of predominantly conjugated bile salts (15). Although relative bile salt proportions following Bulevirtide treatment differed between *Slco1a/1b^−/−^* FVB and *Slco1a/1b^−/−^* B6 models, both profiles were dominated by TCA, CA, and TβMC (Fig. 4C, Fig 5C). These changes were associated with attenuation of DSS-induced colitis in both models. In our previous study, Bulevirtide also elevated plasma TCA and TβMC levels, which was associated with reduced cholestatic liver injury induced by 3,5-diethoxycarbonyl-1,4-dihydrocollidine (DDC), bile duct ligation, or ATP8B1 deficiency in C57BL/6J mice (10). These consistent effects of Bulevirtide on bile salt levels and composition in different colitis and cholestasis studies highlight its potential as a therapeutic strategy for PSC-UC. Additionally, a recent preclinical study suggested that colitis may reduce cholestatic liver damage by suppressing bile salt synthesis (8); however, we did not observe lower plasma bile salt levels in Bulevirtide-treated mice before and after DSS (data not shown).

The beneficial effects of Bulivertide in colitis, were not accompanied by changes in immune cell composition. In the C56BL/6J model, DSS increased colonic immune cell proportions, including monocytes, macrophages, neutrophils and effector memory T cells indicating intestinal inflammation corresponding to previous publications (31, 32). The expansion of macrophages and central effector memory T cells in the mLNs, together with the increased colon weight, is consistent with previous findings in colitis models (33–35). Although RORγt^+^ CD4^+^ T-cells are typically increased in response to DSS (23, 24), they were reduced in the colon in our model, suggesting a limited role in driving colonic inflammation. While Song et al. (36) showed that bile salt supplementation increased the colonic RORγt^+^Helios^-^Treg fraction and alleviated DSS-colitis in mice, we observed a DSS-induced reduction of this Treg population, with Bulevirtide exerting no effect. Nevertheless, in DSS-treated C56BL/6J mice, Bulevirtide induced a less inflammatory colon profile, as indicated by reduced *Tnf* expression, suggesting that its beneficial effects are mediated through modulation of immune cell activation rather than through changes in immune cell composition. This is supported by the observation in LPS-treated *Slco1a/1b^-/-^* FVB mice, where Bulevirtide pre-treatment attenuated inflammation, evidenced by reduced TNF and elevated plasma IL-10 levels. Macrophages are a major source of TNF (37), a primary target of IL-10 (38), and the principal mediators of IL-1β secretion via inflammasome activation (39). In line with the observations in Bulevirtide-treated mice, TCDC dampened the LPS-induced inflammatory response in mouse BMDMs, witnessed by reduced TNF, increased IL-10, as well as suppression of inflammasome-dependent cytokines IL-1β and IL-18. Similar effects were observed in human macrophage BlaER1 cells, as *TNF* and *IL1B* gene expression were gradually reduced by TCDC, most strongly at 200 µM, a concentration reached by Bulevirtide treatment (13). Collectively, our findings suggest that the therapeutic effects of Bulevirtide-mediated bile salt elevation in colitis arise from attenuation of macrophage-driven inflammatory responses.

Other studies report that certain bile salts can activate the inflammasome in macrophages via FXR (20, 40), while we and others show immunosuppressive effects by (conjugated) bile salts in these cells, likely via TGR5 activation (41). The pro-inflammatory effects of bile salts were demonstrated by experiments with hydrophobic, mostly unconjugated bile salts such as CDCA and DCA, which can passively diffuse into cells, thereby promoting inflammation and apoptosis (20, 40, 42, 43). In contrast, Bulevirtide caused a shift in the bile salt pool composition from an unconjugated to a conjugated profile. Conjugated bile salts enter cells only via membrane transport proteins and are less toxic (44). In macrophages, conjugated bile salts are not taken up and do not accumulate, but promote immunomodulatory functions via bile salt-responsive receptors on the cell membrane.

In conclusion, our findings support repurposing of Bulevirtide, a therapeutic peptide for chronic HDV infection, as a future pharmacological therapy for PSC-UC, as NTCP inhibition has hepatoprotective effects in cholestasis and also displays an immune dampening effect in colitis in preclinical settings.

## Methods

### Study approval

This study involved animal experiments that were approved by the Central Authority for Scientific Procedures on Animals (CCD, the Netherlands), license number AVD11400202317642, following positive ethical advice from the local Animal Ethics Committee (DEC).

### Animal experiments

Male and female wild type (WT) C57BL/6J and FVB mice were purchased from Envigo (Venray, the Netherlands). To study hepatic bile salt dynamics mimicking the human situation, male and female *Slco1a/1b* cluster knockout mice (FVB background, hereafter called *Slco1a/1b^-/-^* FVB) were purchased from Taconic (Silkeborg, Denmark) (21). In addition to FVB mice, C57BL/6J WT, *Slco1a1^−/−^* and *Slco1a1/1a4/1b2^−/−^* mice, all on a *Ldlr^−/−^* background, were used. Slco*1a1/1a4/1b2^−/−^* mice (hereafter called *Slco1a/1b^-/-^* B6) were generated using CRISPR-Cas9 as described in Fig.S1. Zygotes isolated from previously generated *Slco1a1^−/−^*/*Ldlr^−/−^*mice (45), were injected with *SpCas9* mRNA and two gRNAs targeting exon 5 of *Slco1a4* and exon 4 of *Slco1b2*. Oligo sequences to generate gRNAs are displayed in Table S1. Embryos were transferred into foster mice, from which offspring was bred homozygously to yield *Slco1a1/1a4/1b2^-/-^/Ldlr^−/−^* mice. Heterozygous *Slco1a1^+/−^/Ldlr^-/-^* mice were intercrossed to generate WT littermates.

For genotyping the *Slco1a4* and *Slco1b2* knockout alleles, polymerase chain reaction (PCR, cat. F-530L, Thermo Fisher, MA, USA) was performed on genomic DNA to amplify the gene regions containing the CRISPR target sites. Primers (Sigma-Aldrich, Grand Island, NY, USA) used are listed in Table S2. The resulting amplicons were cloned into pCR™-TOPO™ vectors (cat. No. K461020, Thermo Fisher). The plasmids were transformed into competent Escherichia coli cells. Since each bacterium takes up only a single plasmid, this step allowed the separation of individual alleles that may have arisen from CRISPR-Cas9 editing. Plasmid DNA was then isolated and sequenced by BigDye Terminator sequencing according to the manufacturer’s instructions (cat. No. 4337450, Thermo Fisher). Sequencing revealed three point mutations in *Slco1a4* that resulted in two amino acid substitutions, whereas a 7-nucleotide deletion in *Slco1b2* caused a frameshift and consequent introduction of a premature stop codon (Fig. S1B).

All mice were housed at the Amsterdam University Medical Center, Amsterdam under specific pathogen-free conditions in ventilated cages. The mice were maintained on a 12-h light/dark cycle and provided with water and chow *ad libitum*.

Effects of inhibition of hepatic bile salt uptake on LPS-induced acute inflammatory responses *in vivo* were assessed in male *Slco1a/1b*^-/-^ mice, which were administered a single dose of Bulevirtide (2.5 µg/g BW) or vehicle and cholecystokinin (cat. No. ab120209, Abcam, Cambridge, England) via the tail vein. Bulevirtide was dissolved in a vehicle containing 25 mM sodium carbonate (cat. No. 6392, Merck, Burlington, MA, USA) at pH 8.8 and 50 mg/mL mannitol (cat. No. M4125, Sigma-Alrich). One hour after administration of Bulevirtide or vehicle, LPS (cat. No. tlrl-eblps, Invivogen, San Diego, CA, USA) was administered by intraperitoneal injection to the mice (20 µg/g BW).

Colitis was induced by drinking water supplemented with DSS (cat. No. DB001-500g, TdB consultancy, Uppsala, Sweden) for 5 days. Female WT and *Slco1a/1b^-/-^*FVB mice received 6% (w/v) DSS. Male *Slco1a/1b^-/-^* B6 received 2.5% DSS. Simultaneously, mice received daily subcutaneous injections with Bulevirtide (2.5 µg/g BW) or vehicle. Seven to eight days after the start of DSS supplementation, animals were euthanized.

Daily water intake was determined per cage, which housed 2 to 6 mice. Macroscopic disease score was determined by assessing the presence of diarrhea (score 0-3), inflammation (0–3) and edema in the colon (0–3). Histological disease assessment was scored according to the standardized scoring system described previously (46). All scoring was performed by blinded observers.

To examine plasma clearance and biliary excretion of bile salts in C57BL/6J strains, WT, *Slco1a1^−/−^*, and *Slco1a/1b^-/-^* B6 mice were treated with Bulivertide (2.5 µg/g BW) or vehicle to inhibit NTCP at Zeitgeber Time (ZT)4 (i.e. 4 hours into the light phase). After 2 hours, mice were anesthetized with ketamine/xylazine (120 and 10 mg/kg, respectively). The gallbladder was cannulated, and the common bile duct was distally ligated. A second cannula was inserted in the vena jugularis for intravenous injection. Next, 5 mL/kg BW 30 mM sodium taurocholic acid (TCA) (T4009, Sigma-Aldrich, Saint Louis, Missouri, USA; dissolved in 0.9% NaCl) with 75 μCi/kg BW [^3^H]TCA (NET322, PerkinElmer) was infused via the vena jugularis. Blood and bile were sampled via a tail vein cut and a gallbladder cannula, respectively, for up to 60 minutes post-infusion. Plasma (5 μl) or bile (total collected volume) was directly added to 2.5 ml Ultima Gold, and radioactivity was measured using a liquid scintillation counter (Tri-Carb 2900 TR, PerkinElmer). ^3^H[TCA] activity in plasma or bile was expressed as a percentage of the injected dose.

Animal studies were performed in accordance with the ARRIVE 2.0 guidelines. Sample size was determined by a priori power calculations based on the primary outcome measures, including cytokine concentrations (LPS-induced inflammation) and disease severity (colitis), to ensure sufficient statistical power across experiments. Animals were randomized to treatment groups. Treatment and sacrifice were carried out in a standardized order to minimize potential confounding. All animals were included in the statistical analyses.

### Sex as a biological variables

Male FVB mice were used for the systemic LPS model to obtain robust and reproducible pro-inflammatory cytokine responses (47). Male C57BL/6 mice were included in the DSS model because this strain–sex combination demonstrates well-established susceptibility to colitis (22). DSS colitis was assessed in female FVB mice to incorporate other sex and strain as biological variables.

### Isolation of leukocytes from the colon

Isolation of colon leukocytes was performed as described previously (48). Post-isolation, colons were placed on a paper towel soaked with phosphate-buffered saline (PBS), opened longitudinally and cleaned by scraping the intestinal content with a spatula. Cleaned colons were washed in Hanks’ Balanced Salt Solution (HBSS, without calcium/magnesium, Thermo Fisher) supplemented with 2 mM ethylenediaminetetraacetic acid (EDTA, Thermo Fisher), cut into small pieces and stored in 10 ml HBSS on ice. After sample collection, colon sections were strained using 250 µM Nitex and washed by incubating for 10 minutes in 10 ml HBSS/EDTA at 37 °C under agitation (200 RPM) to remove mucous and strip epithelial cells. After 10 minutes, samples were strained through 250 µM Nitex and washed 2 additional times as described. Post-washing, colon samples in the FVB model were digested in HBSS supplemented with 2% fecal calf serum (FCS), 60 μg/mL Liberase TL (Cat. No. 05401119001, Roche, Basel, Switzerland), and 40 μg/mL DNAse (cat. No. 05401020001 and 11284932001, Sigma-Aldrich) for 40 minutes while stirring. In the C57BL/6J model, colon samples were digested for 30 minutes at 37°C under agitation (200 RPM) in 5 ml digestion mix consisting of RPMI 1640 + Glutamax supplemented with 1 mg/ml Collagenase Type IV from *Clostridium histolyticum* (Sigma-Aldrich, 125 CDU/ml), 1 mg/ml Dispase II (Sigma-Aldrich, 1.4 U/ml), 1 mg/ml Collagenase D from *C. histolyticum* (Roche, 250 Mandl U/ml) and DNase I (Sigma-Aldrich, 2000 U/ml). After digestion, samples were strained (100 µM strainer, Corning, NY, USA), washed with PBS supplemented with 0.5% bovine serum albumin (BSA) and 2.5mM EDTA and pelleted at 450 x g for 10 minutes at 4°C. Samples were subsequently strained (40 µM strainer, Corning), washed in PBS/BSA/EDTA, and 1*10^6^ cells per sample were processed for flow cytometry (see subsection flow cytometry).

### Isolation of leukocytes from mesenteric lymph node and spleen

Spleen and mesenteric lymph node samples were collected in RPMI 1640+Glutamax and stored on ice. After sample collection, the samples were ruptured using a 1 ml syringe and subsequently digested in RPMI 1640 + glutamax supplemented with 1 mg/ml Collagenase D from *C. histolyticum* (Roche, 250 Mandl U/ml) and DNase I (Sigma-Aldrich, 2000 U/ml) for 20 minutes at 37°C. After digestion, samples were strained (100 µM strainer, Corning), pelleted at 450 x g for 5 minutes at 4°C and washed with PBS/BSA/EDTA. Next, cell pellets were treated with erythrocyte lysis buffer consisting of 0.15 M NH_4_Cl (Merck), 1mM KHCO_3_ (Merck) and 0.1 mM EDTA (Invitrogen, Waltham, MA, USA) for 2 minutes at room temperature and subsequently washed with PBS/BSA/EDTA. After washing, samples were counted and 1*10^6^ cells per sample were further processed for flow cytometry (see subsection flow cytometry).

### Flow cytometry

In the FVB model, macrophage polarization was assessed. Gating strategies are shown in Fig. S3D. Antibodies used were obtained from Biolegend (San Diego, CA, USA): αCD45-APC-Cy7 (clone 30-F11), αCD11b-PerCP (clone M1/70), αCD64-PE (clone X54–5/7.1), αCD206-AF488 (clone MR5D3), from BD Biosciences (San Jose, CA, USA): αLy6G-AF700 (clone 1A8), and from eBioscience (San Diego, CA, USA): αLy6C-APC (clone HK1.4). Cells were stained using 4’,6-diamidino-2-fenylindool (DAPI) to discriminate against live cells. Cells were analyzed using a FACS Fortessa (BD Biosciences) and analyzed using FlowJo software (Treestar Inc., San Jose, CA, USA). Colonic macrophages were defined as DAPI^−^CD45^+^CD11b^+^Ly6G^-^CD64^+^.

In the C57BL/6J model, two different panels were used to examine immune populations; representative gating strategies can be found in Fig. S5. Samples were processed and acquired as described previously (49). A full list of reagents and their description can be found in Table S3. Briefly, isolated samples were first incubated with Zombie-NIR fixable viability dye (Biolegend, San Diego, CA, USA) in PBS supplemented with True Stain Monocyte Blocker (Biolegend) and FC-block (Biolegend) for 20 minutes at room temperature. Subsequent live staining of specific targets prior to fixation (see Table S3) was performed in PBS/BSA/EDTA for 30 minutes on ice. Fixation was performed using the eBioscience Foxp3/Transcription factor staining buffer set (Invitrogen). Post-fixation, the remaining cell surface targets were stained in PBS/BSA/EDTA supplemented with Brilliant Stain Buffer Plus (BD

Biosciences) and True Stain Monocyte Blocker (Biolegend) for 30 minutes at 4°C. After staining, the samples were washed twice using permeabilization buffer (Invitrogen), and the intracellular targets (see Table S3) were stained in permeabilization buffer supplemented with Brilliant Stain Buffer Plus (BD Biosciences and True Stain Monocyte Blocker (Biolegend) for 30 minutes at 4°C. After staining, all samples were washed twice with PBS/BSA/EDTA and acquired on a Cytek Aurora 5-laser spectral flow cytometer (Cytek Biosciences, Fremont, CA, USA). Acquired samples were unmixed using SpectroFlow v3 (Cytek Biosciences) and the data were analyzed using FlowJo v10.10 (BD Biosciences).

### Cytokine measurements

Cytokines TNF, IL-10, IL-6, macrophage attractant protein-1 (MCP1) and IL-12p70 in mouse plasma were measured using the BD^TM^ CBA mouse inflammation kit according to the manufacturer’s instructions (cat. No. 552364, BD Biosciences). Measurements were performed on a FACS Fortessa (BD Biosciences) and analyzed using FlowJo software (Treestar Inc.). IL-10, IL-1β and TNF concentrations were measured in BMDM supernatant by ELISA (cat. No. DY401, DY417, DY410, R&D Systems, Minneapolis, MN, USA) according to the manufacturer’s instruction.

### Bile acid measurements

Concentrations of individual bile acid species in plasma were measured with HPLC as described previously (50).

### Histology

Colon processing for histological assessment was performed as described previously (46). 5 µm slides were cut and stained with hematoxylin and eosin (H&E) as described before (26).

### Cell culture

Bone marrow-derived macrophages (BMDMs) of WT C57BL/6J mice were differentiated as described previously (51). U2OS cells (human bone osteosarcoma epithelial cells) and U2OS cells stably expressing NTCP (52) were cultured in DMEM supplemented with 2 mM L-glutamine (Lonza), 100 U/L Penicillin/Streptomycin (Lonza) and 10% FCS (Gibco). Recently developed human transdifferentiation cell culture system BlaER1 which can develop from malignant B cells to macrophages (53) were cultured in RPMI 1640 Medium without Phenol Red supplemented with 2 mM L-Glutamine (Lonza), 1 mM Sodium Pyruvate (Lonza), 100 U/mL Penicillin, 100 μg/mL Streptomycin (Lonza), and 10% charcoal-treated FCS (Gibco). For transdifferentiation into macrophages, cells were transferred into medium containing 10 ng/mL human IL-3, 10 ng/mL human macrophage colony-stimulating factor (M-CSF) (Miltenyi Biotec), and 100 nM β-Estradiol (Sigma-Aldrich). After differentiation, both BMDMs and BlaER1 cells were treated with taurochenodeoxycholic acid (TCDC) (T6260, Sigma-Aldrich) for 1 hour, before stimulation with LPS (cat. No. L4130, Sigma-Aldrich) 10 ng/ml for 2.5 hours. Then, 1 mM adenosine triphosphate (ATP) (cat. No. A2383, Sigma-Aldrich) was added during the final 30 minutes before harvesting. In 12-well plate for qPCR analysis, 1×10^6^ BlaER1 cells were plated; while 7.5×10^5^ BMDMs were plated in 12-well plates for ELISA and qPCR experiments. For protein quantification, 2×10^6^ BMDMs were plated in 6 well plates.

### Bile salt uptake assay

Bile salt uptake activity was assessed in BMDMs, U2OS parental and U2OS cells stably expressing NTCP cells as previously described (54). Briefly, cells were washed twice with warm uptake buffer (5 mM KCl, 1.1 mM KH_2_ PO_4_, 1 mM MgCl_2_, 1.8 mM CaCl_2_, 10 mM D-glucose, 10 mM Hepes, and 136 mM NaCl). Then, cells were incubated at 37 °C for 2 minutes with uptake buffer containing 20 μM taurocholic acid (TCA) (T4009, Sigma) of which a trace amount was tritium-labelled (^[3H]^TCA; Perkin Elmer, Groningen, the Netherlands). Finally, cells were washed four times with ice-cold PBS buffer and lysed with milliQ water containing 0.05% (w/v) SDS. Radioactivity in the lysates was measured by liquid scintillation counting.

### Caspase assay

In 96-well plates, 4×10^4^ BMDMs were plated and treated for 3 hours with different concentrations of TCDC. Cells were cultured in RPMI medium. Raptinal (20 µM) was used as a positive control for apoptosis induction. Caspase 3/7 activity was measured with an artificial peptide substrate (Ac-DEVD)2-Rh110 using the SensoLyte Homogeneous Rh110 Caspase 3/7 Assay Kit (AnaSpec Inc., Fremont, CA, USA) according to the manufacturer’s instructions. The kinetics of fluorophore Rh110 release, as a result of cleavage by caspase 3 and caspase 7, was measured by NOVOstar microplate reader (BMG Labtech GmbH, Offenburg, Germany) at λex / λem = 480 / 510 nm. The slope in the linear window of the kinetic reading was calculated and defined as the caspase 3/7 activity.

### Cell viability assay

Cell viability was measured by using Roche Cell Proliferation Reagent WST-1 (Sigma-Aldrich) according to the manufacturer’s protocol and spectrophotometrically quantified using the Synergy HTX Multi-Mode reader (BioTek/Agilent).

### RNA isolation, cDNA synthesis and qPCR

Total RNA isolation and qPCR was performed as described previously (15). Colonic RNA was isolated with the Isolate II RNA Mini Kit (Bioline, Cincinnati, OH, USA). Primers (Sigma-Aldrich) used are listed in Table S4. Relative gene expression was determined by qPCR. Results were processed using the BioRad CFX Maestro 5.0 software and LinRegPCR 12.5 software, and expression levels were normalized to two reference genes, 36B4 and HRPT.

### Western blot

Whole cell lysates were obtained as described previously (15). Proteins were transferred by wet-blotting to PVDF-membrane and probed with anti-IL-1β (cat. No. AF-401-NA, R&D Systems), anti-caspase-1 (cat. No. AG-20B-0042-C100, AdipoGen, San Diego, CA, USA) or anti-GAPDH (cat. No. CST 2118S, Cell signaling/Bioke, Danvers, MA, USA). Immune complexes were detected with a horseradish peroxidase-conjugated secondary antibody (Biorad, CA, USA), visualized using enhanced chemiluminescence detection reagent (Lumi-light, Roche) and detected using ImageQuant LAS 4000 (GE Healthcare, IL, USA).

### Statistics

Data are provided as the mean ± standard deviation. Normal distribution was examined with D’Agostino & Pearson test. Differences between 2 groups were analyzed using Mann-Whitney U test. Group differences were assessed using Kruskal–Wallis test followed by Dunn’s multiple comparisons test with Benjamini–Krieger–Yekutieli false discovery rate (FDR) correction. Statistical significance was considered at *p* < 0.05. Data were analyzed using GraphPad Prism 10.2.0 (GraphPad Software Inc, La Jolla, CA).

### Access to data

All authors had access to the study data and had reviewed and approved the final manuscript. All data and methods supporting the findings of this study are available within the paper and its supplemental materials. All the raw data files will be made available upon reasonable request.

## Abbreviations

ATP: adenosine triphosphate
BMDM: bone marrow-derived macrophages
CA: cholic acid
CXCL10: C-X-C motif chemokine ligand 10
DCA: deoxycholic acid
DDC: 5-diethoxycarbonyl-1,4-dihydrocollidine
DSS: dextran sodium sulfate
FXR: nuclear receptor Farnesoid X receptor
ISG15: interferon-stimulated gene 15
mLNs: mesenteric lymph nodes
MX1: myxovirus resistance protein 1
NLRP3: NLR family pyrin domain containing 3
NFκB: nuclear factor kappa-light-chain-enhancer of activated B cells
NTCP: Na⁺ taurocholate co-transporting polypeptide
OAS1: 2′-5′-oligoadenylate synthetase 1
OCA: obeticholic acid
OATP: Organic Anion Transporting Polypeptide
PBC: primary biliary cholangitis
PSC: primary sclerosing cholangitis
TCA: taurocholic acid
TβMC: tauro-β-muricholic acid
TCDC: taurochenodeoxycholic acid
TNBS: trinitrobenzenesulfonic acid
TGR5: Takeda G-protein-coupled receptor 5
UDCA: ursodeoxycholic acid.

## Author contributions

Concept and design of the experiments: TAN, RLRA, PJK, WIhP, MEW, CCP, SFJvdG. Experiments and procedure: TAN, RLRA, PJK, JML, WIhP, DRW, IB, SD, EV. Writing – original draft: TAN, RLRA, CCP, SFJvdG. Writing – review and editing: all authors.

## Data transparency

All data and methods supporting the findings of this study are available within the paper and its supplemental materials. All the raw data files will be made available upon reasonable request.

## Acknowledgments

The authors thank Claire Groenen and Dandan Wu for their assistance with immune cell isolation for flow cytometry.

