## Supplemental figures and tables for "Increasing plasma bile salt levels with Bulevirtide alleviates DSS-induced colitis and LPS-induced inflammation"

**Supplemental materials**

**Supplemental figures**

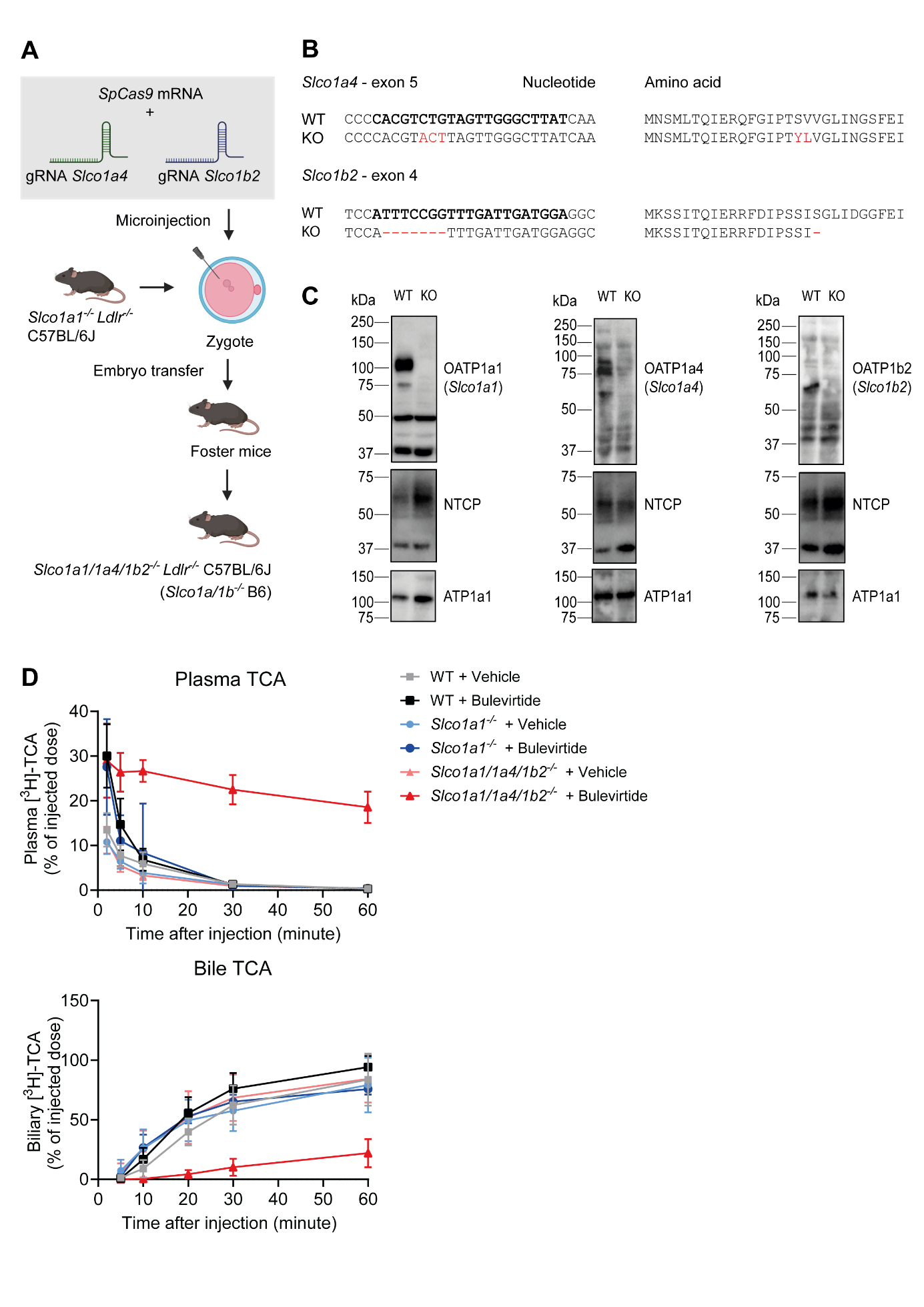

**Fig. S1. Generation and characterization of the *Slco1a1/1a4/1b2^-/-^/Ldlr^-/-^* C57BL/6J (*Slco1a/1b^-/-^* B6) mouse model.** (A) Zygotes from *Slco1a1^−/−^/Ldlr^−/−^* mice were injected with *SpCas9* mRNA and two gRNAs targeting exon 5 of *Slco1a4* and exon 4 of *Slco1b2.* Embryos were implanted in foster mice, from which offsprings were bred homozygously into *Slco1a1/1a4/1b2^-/-^/Ldlr^-/-^* (*Slco1a/1b^-/-^* B6)*.* (B) Alignment of wild*−*type (WT) and knockout (KO) alleles shown for nucleotide and predicted amino acid sequences. Modifications introduced at the gRNA binding sites—indicated in bold—are highlighted in red. Genotyping was performed by Sanger sequencing. (C) Hepatic abundance of OATP1A1, OATP1A4, and OATP1B2 measured in 3 representative offspring. (D) Plasma clearance and biliary excretion of [^3^H]TCA in WT, *Slco1a1^−/−^ and Slco1a1/1a4/1b2^-/-^* (n=4-8/group). Mice were injected with vehicle or Bulevirtide (2.5 µg/g) BW. The gallbladder was cannulated, and the common bile duct was ligated for bile collection. 5 mL/kg BW 30 mM TCA with 75 μCi/kg BW [^3^H]TCA was infused via the vena jugularis, and plasma and bile were sampled up to 60 minutes post-infusion to measure [^3^H] activity.

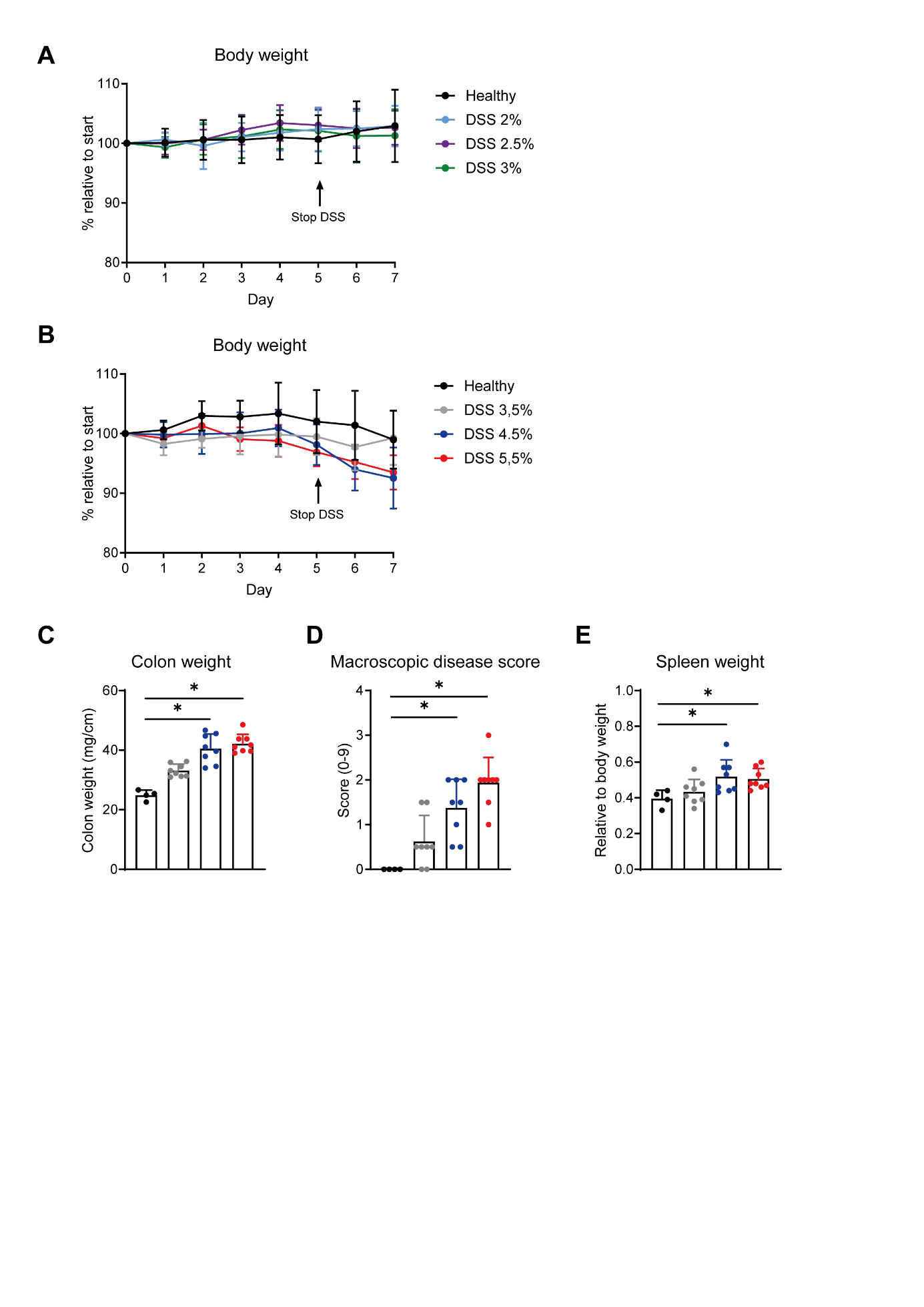

**Fig. S2. A high dose of DSS supplementation is required to induce colitis in FVB mice.** DSS concentrations needed to induce colitis in FVB WT mice were determined by supplementing DSS in drinking water for 5 days. Body weight relative to start of DSS in mice receiving low-dose DSS (2-3%) (A) and high-dose DSS (3.5-5.5%) (B). Colon weight (C), macroscopic disease score (D), and spleen weight (E) at sacrifice of mice receiving high dose DSS. Statistical significance was assessed with Kruskal-Wallis test in (C, D, E). * *p<*0.05.

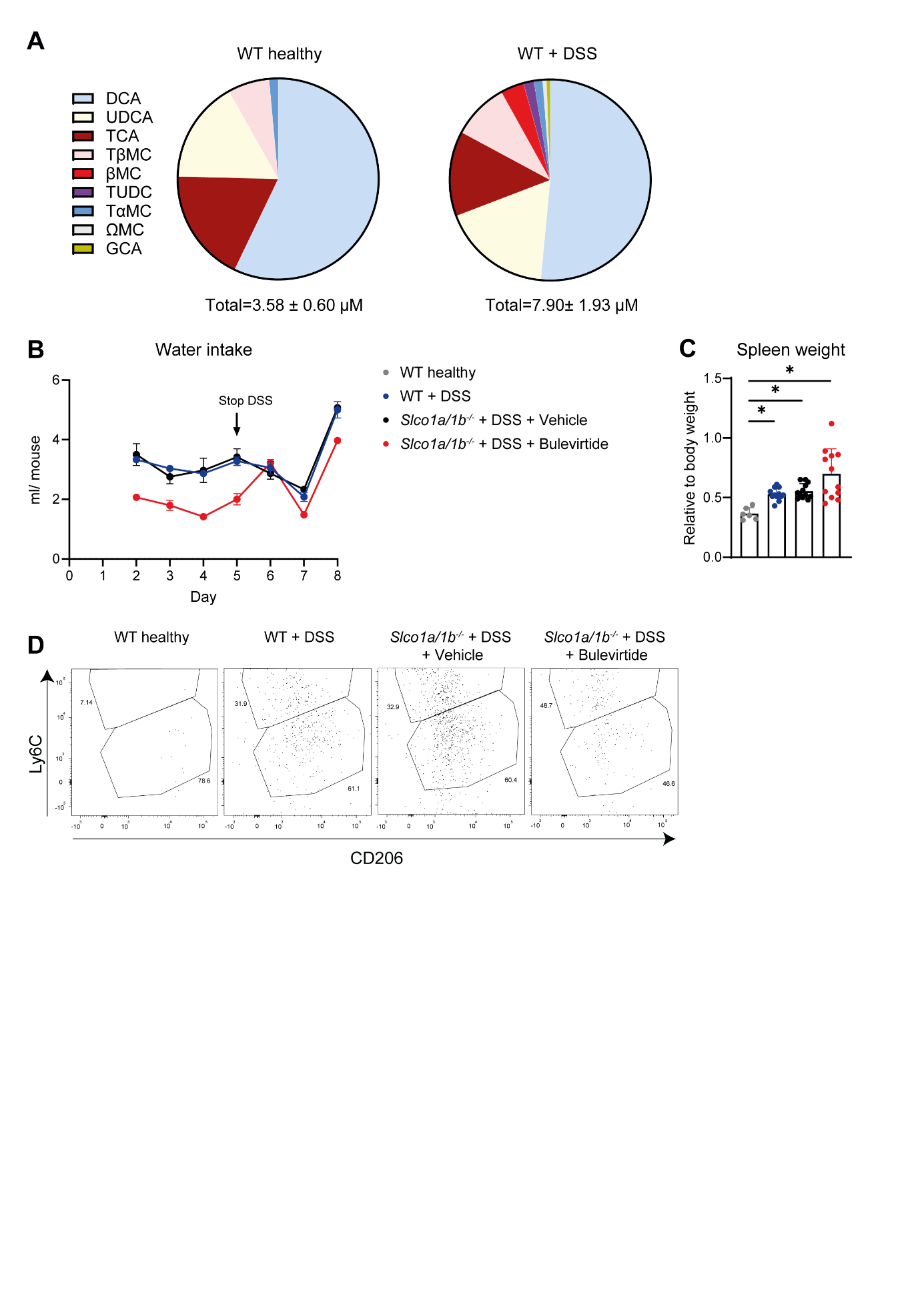

**Fig. S3. Additional data from the colitis study in FVB mice.** (A) Concentration of individual bile salt species in WT FVB and WT FVB treated with DSS mice, measured by HPLC. (B) Water intake per mouse; DSS supplementation was started on day 1 and stopped on day 5. (C) Spleen weight at sacrifice. (D) Gating strategy for flow cytometry of colon; Myeloid cells were gated as DAPI^−^CD45^+^Ly6G^−^CD11b^+^CD64^+^ cells and subdivided into regulatory (CD206^hi^Ly6C^lo^) and inflammatory (CD206^lo^Ly6C^hi^) macrophages.

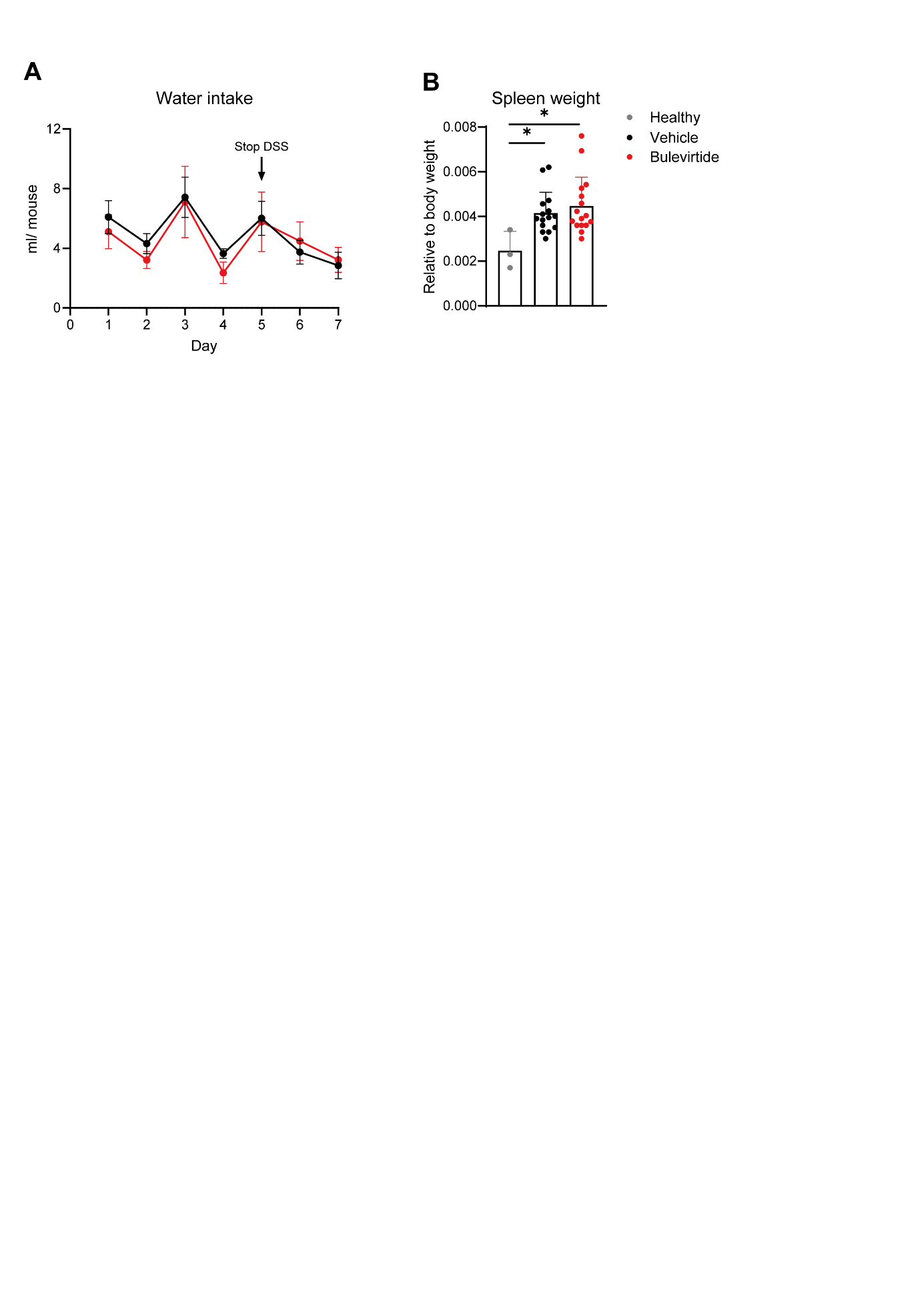

**Fig. S4. Additional data from the colitis study in *Slco1a/1b*^-/-^ B6** **mice.** (A) Water intake per mouse; DSS supplementation was started on day 1 and stopped on day 5. (B) Spleen weight at sacrifice. Data are means ± SD. Statistical significance was assessed with Kruskal-Wallis test in (B). * *p<*0.05.

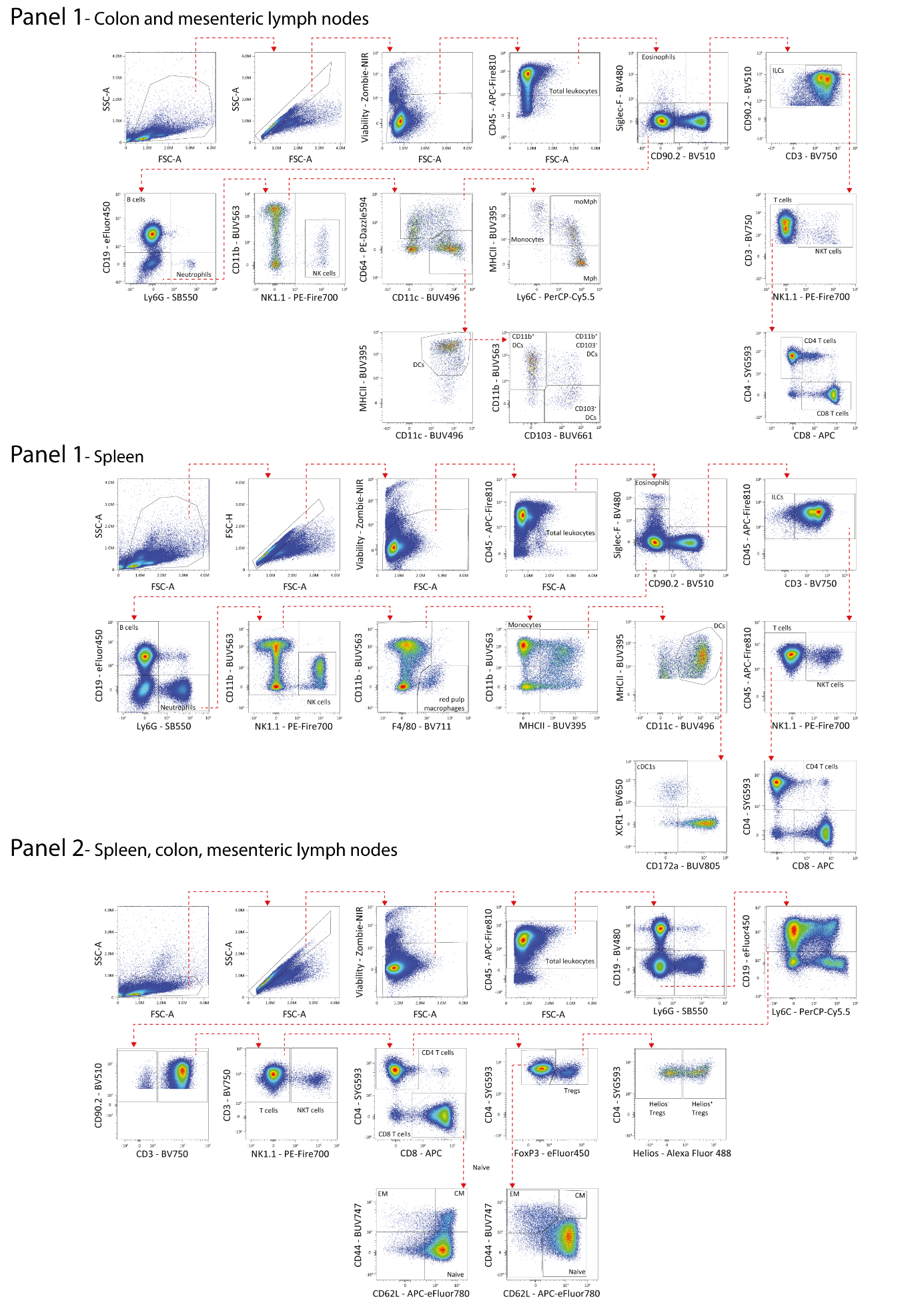
**Fig. S5. Gating strategies of flow cytometry in the colitis study in *Slco1a/1b*^-/-^ B6** **mice.** Representative gating strategies used to identify described immune populations in the spleen, colon and mesenteric lymph node. For more detailed information on antibodies and panels please refer to Table S3.

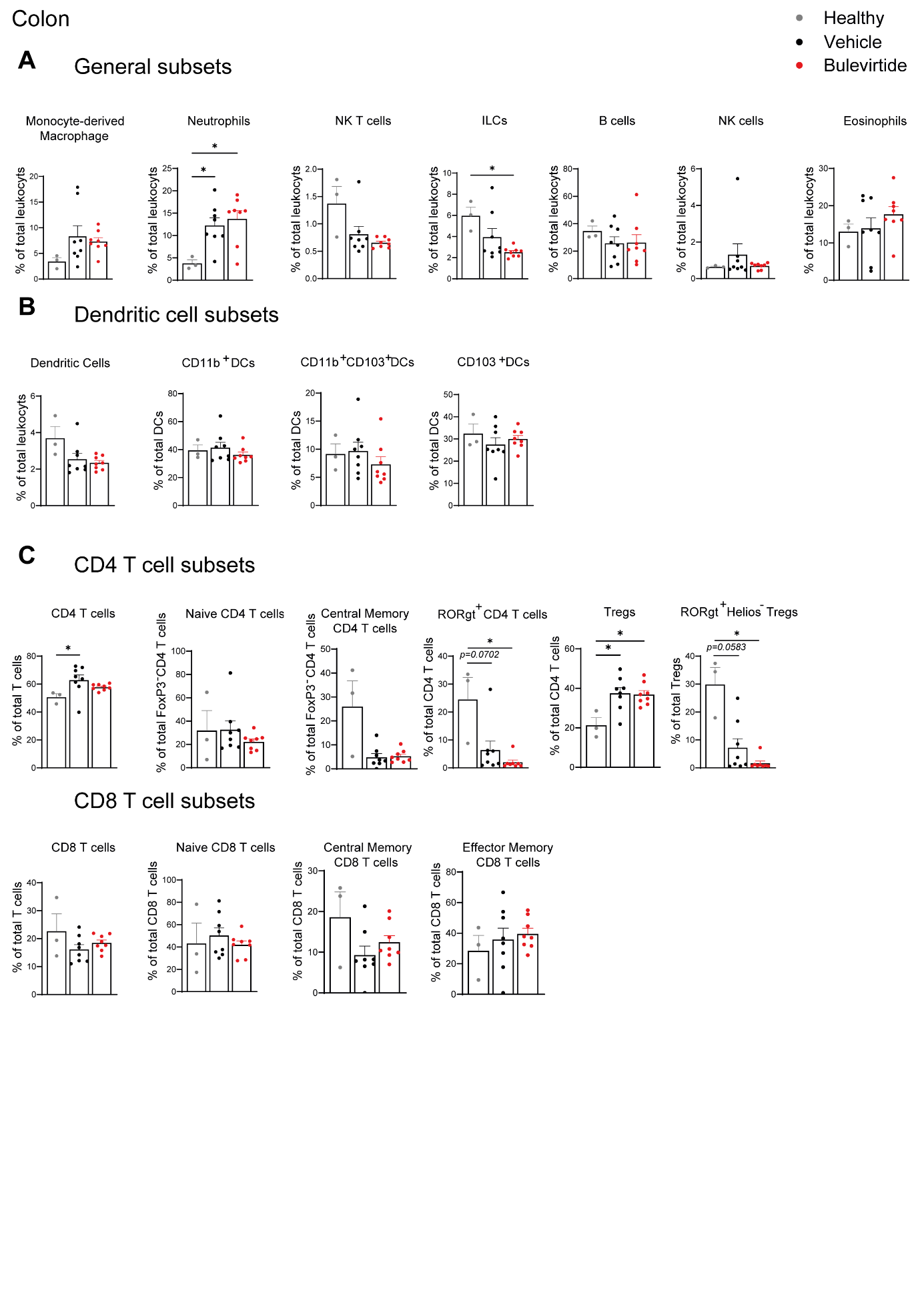

**Fig. S6. Immune cells population in colon in the colitis study in *Slco1a/1b*^-/-^ B6** **mice.** (A) Plots displaying major immune cell subsets in the colon as frequency of the total leukocyte pool. (B) Total dendritic cells (DCs) and DC subsets plotted as frequency of total leukocytes and total DCs respectively. (C) CD4, CD8 T cells presented as frequency of total T cells; naïve, effector memory and central memory CD4 and CD8 T cells displayed as frequency of total CD4 and total CD8 T cells respectively; regulatory T cells (Tregs) and RORgt^+^ CD4 T cells presented as frequency of total CD4 T cells; RORgt^+^ Helios^-^ Tregs displayed as frequency of total Tregs. Data are means ± SD. Statistical significance was assessed with Kruskal-Wallis test. * *p<*0.05.

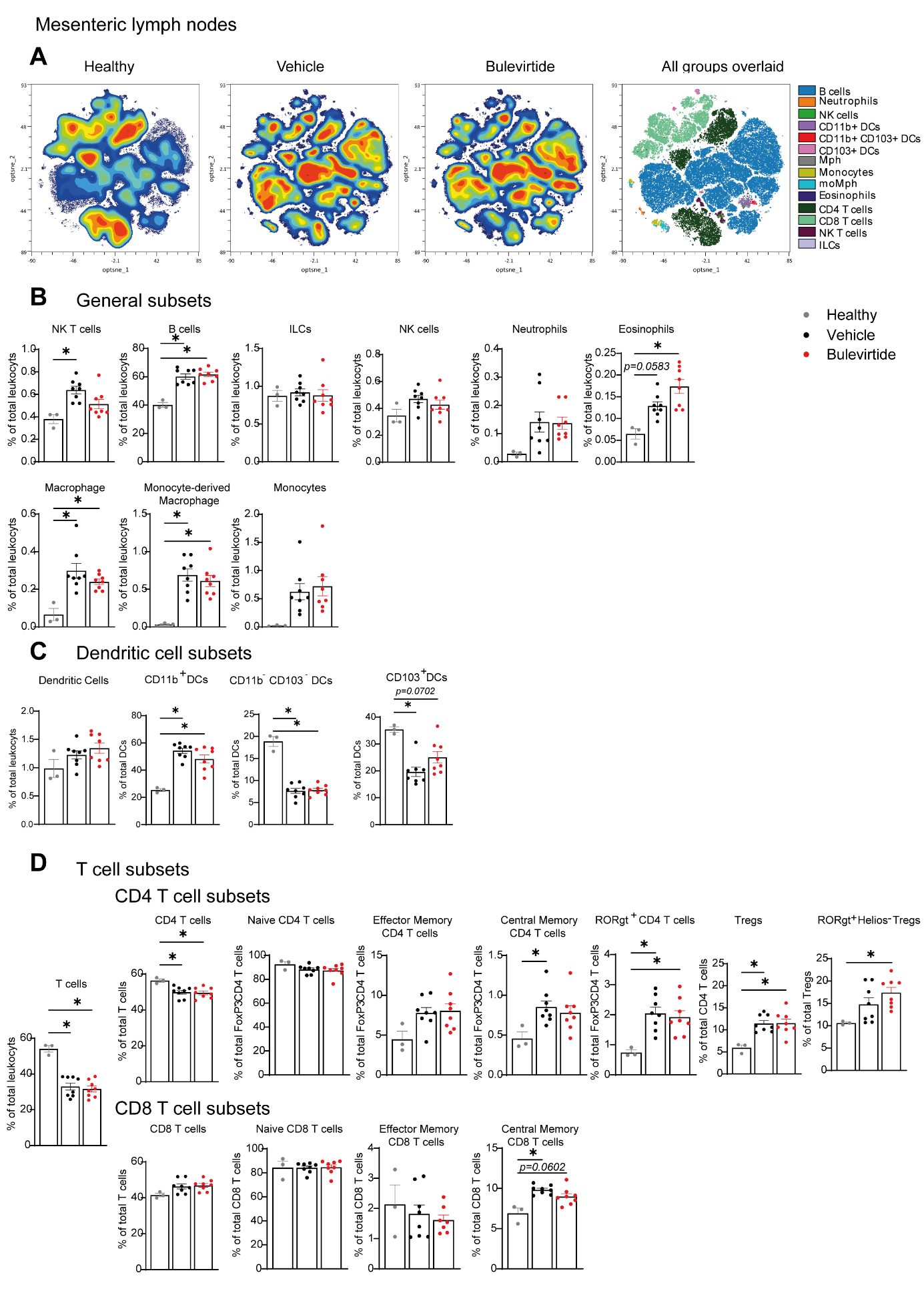
**Fig. S7. Immune composition of the mesenteric lymph nodes in the colitis study in *Slco1a/1b*^-/-^ B6** **mice.** (A) Optimized t-Distributed Stochastic Neighbor Embedding (optsne) used to visualize global changes in immune composition of the mesenteric lymph node in healthy, vehicle and Bulevirtide treated samples with each plot showing pooled data from all mice per group. (B) Plots displaying major immune cell subsets in the mesenteric lymph node as frequency of the total leukocyte pool. (C) Total dendritic cells (DCs) and DC subsets plotted as frequency of total leukocytes and total DCs respectively. (D) T cells and CD4 and CD8 T cells presented as frequency of total leukocytes and total T cells respectively; naïve, effector memory and central memory CD4 and CD8 T cells shown as frequency of total CD4 and CD8 T cells respectively; RORgt^+^ CD4 T cells and regulatory T cells plotted as frequency of total CD4 T cells; RORgt^+^ Helios^-^ Tregs displayed as frequency of total Tregs. Data are means ± SD. Statistical significance was assessed with Kruskal-Wallis test. * *p<*0.05.

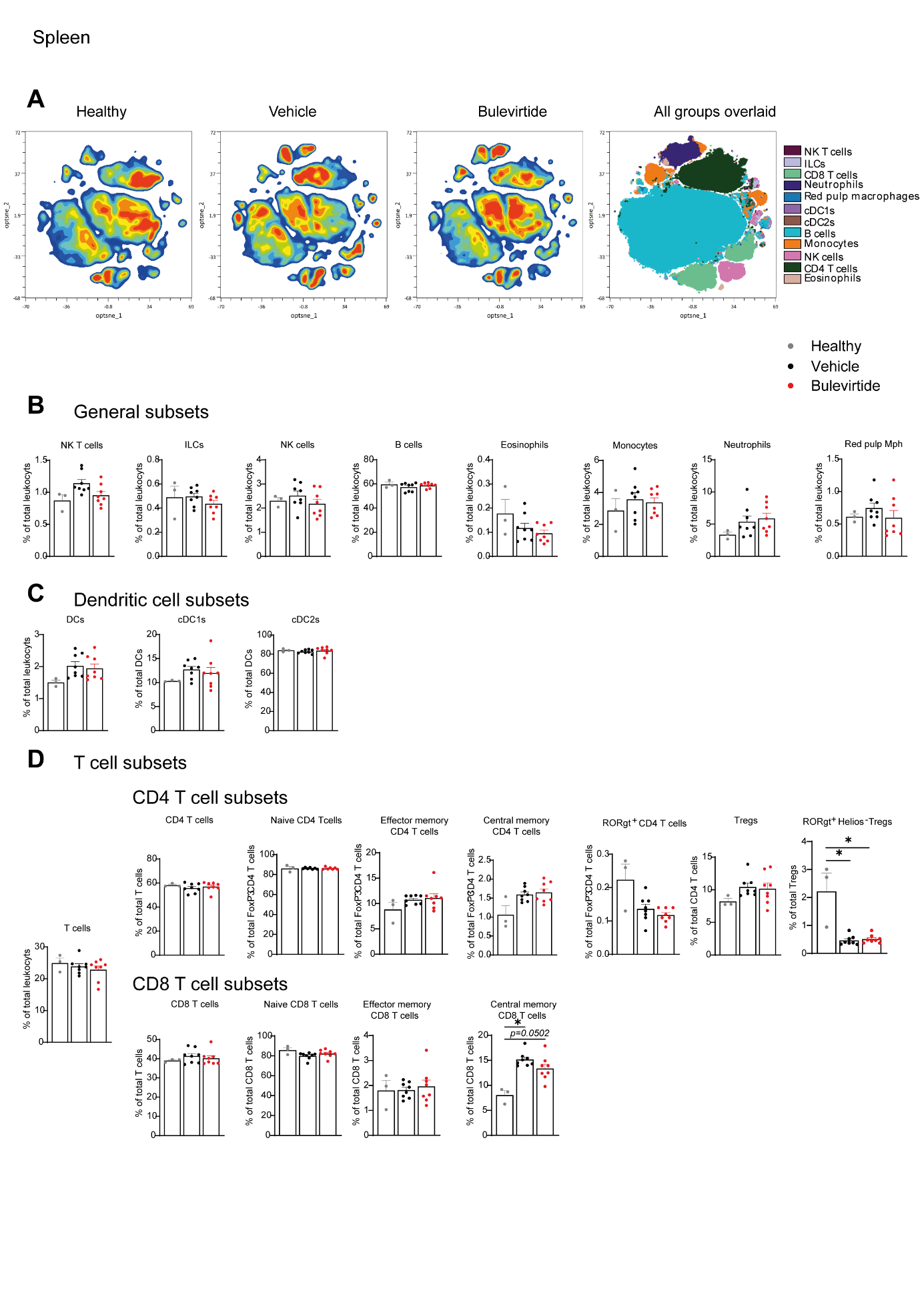

**Fig. S8. Immune cells population in spleen in the colitis study in *Slco1a/1b*^-/-^ B6** **mice.** (A) Optimized t-Distributed Stochastic Neighbor Embedding (optsne) used to visualize global changes in immune composition of the spleen in healthy, vehicle and Bulevirtide treated samples with each plot showing pooled data from all mice per group. (B) Plots displaying major immune cell subsets in the spleen as frequency of the total leukocyte pool. (C) Total dendritic cells (DCs) and DC subsets plotted as frequency of total leukocytes and total DCs respectively. (D) T cells and CD4 and CD8 T cells presented as frequency of total leukocytes and total T cells respectively; naïve, effector memory and central memory CD4 and CD8 T cells shown as frequency of total CD4 and CD8 T cells respectively; RORgt^+^ CD4 T cells and regulatory T cells plotted as frequency of total CD4 T cells; RORgt^+^ Helios^-^ Tregs displayed as frequency of total Tregs. Data are means ± SD. Statistical significance was assessed with Kruskal-Wallis test. * *p<*0.05.

**Supplemental tables**

**Table S1. mSlco1a4 and mSlco1b2 oligo’s for sgRNA**

| exon 5 of *Slco1a4* | 5’ −TAGG[ATAAGCCCAACTACAGACGT](https://design.synthego.com/) − 3’  3’ − TATTCGGGTTGATGTCTGCACAAA − 5’ |
| --- | --- |
| exon 4 of *Slco1b2* | 5’ −TAGGTCCATCAATCAAACCGGAAA −3’  3’ – AGGTAGTTAGTTTGGCCTTTCAAA −5’ |

**Table S2. Primers used for genotyping**

| Gene | Primers |
| --- | --- |
| exon 5 of *Slco1a4* | Fw: CTAGGCATTTCTGTTGGCATTAAC  Rv: CATGGATCTCAGTCAGATTAAATGATC |
| exon 4 of *Slco1b2* | Fw: GGCAGCCCTCTCATTCAGCTACATATG  Rv: GACAATAAAAATGATATTGCATGTTG |

**Table S3.** Staining reagents of flow cytometry

| Target | Fluorochrome | Clone | Supplier | Catalog# | Pre-fixation staining | Intracellular | Panel |
| --- | --- | --- | --- | --- | --- | --- | --- |
| CD3 | BV750 | 17A2 | Biolegend | 100249 | No | No | 1+2 |
| CD4 | SparkYellowGreen 593 | GK1.5 | Biolegend | 100488 | No | No | 1+2 |
| CD8 | APC | 53-6.7 | TONBO biosciences | 20-0081 | No | No | 1+2 |
| CD11b | BUV563 | M1/70 | BD Biosciences | 741357 | No | No | 1+2 |
| CD11c | BUV496 | N418 | BD Biosciences | 750450 | No | No | 1+2 |
| CD19 | BV480 | 1D3 | BD Biosciences | 566107 | No | No | 2 |
| CD19 | eFluor450 | 1D3 | Invitrogen | 48-0193-82 | No | No | 1 |
| CD44 | BUV737 | IM7 | BD Biosciences | 612799 | Yes | No | 2 |
| CD45 | APC-Fire810 | 30-F11 | Biolegend | 103174 | No | No | 1+2 |
| CD62L | APC-eFluor780 | MEL-14 | Invitrogen | 47-0621-82 | Yes | No | 2 |
| CD64 | PE-Dazzle594 | X54-5/7.1 | Biolegend | 139319 | No | No | 1 |
| CD90.2 | BV510 | 30-H12 | Biolegend | 105335 | No | No | 1+2 |
| CD103 | BUV661 | 2E7 | BD Biosciences | 750718 | Yes | No | 1+2 |
| CD172a | BUV805 | P84 | BD Biosciences | 741997 | No | No | 1+2 |
| F4/80 | BV711 | BM8 | Biolegend | 123147 | No | No | 1+2 |
| FoxP3 | eFluor450 | FJK-16s | Invitrogen | 48-5773-82 | No | Yes | 2 |
| Ly6C | PerCP-Cy5.5 | HK1.4 | Biolegend | 128012 | No | No | 1+2 |
| Ly6G | SparkBlue550 | 1A8 | Biolegend | 127664 | No | No | 1+2 |
| MHCII | BUV395 | 2G9 | BD Biosciences | 743876 | No | No | 1+2 |
| NK1.1 | PE-Fire700 | PK136 | Biolegend | 108774 | No | No | 1+2 |
| RORγt | PE | Q31-378 | BD Biosciences | 562607 | No | Yes | 2 |
| Siglec-F | BV480 | E50-2440 | BD Biosciences | 746668 | No | No | 1 |
| XCR1 | BV650 | ZET | Biolegend | 14822 | Yes | No | 1+2 |

**Table S4. Primers used for qRT-PCR**

| Gene | Species | Primers |
| --- | --- | --- |
| Tnf | mouse | Fw: CATCTTCTCAAAATTCGAGTGACAA  Rv: TGGGAGTAGACAAGGTACAACCC |
| Mcp1 | mouse | Fw: CTTCTGGGCCTGCTGTTCA  Rv: CCAGCCTACTCATTGGGATCA |
| Il1β | mouse | Fw: AAAGAATCTATACCTGTCCTGTGTAATGAAA  Rv: GGTATTGCTTGGGATCCACACT |
| Il10 | mouse | Fw: TTTGAATTCCCTGGGTGAGAA  Rv: CTCCACTGCCTTGCTCTTATTTTC |
| Il12 | mouse | Fw: GGTGCAAAGAAACATGGACTTG  Rv: CACATGTCACTGCCCGAGAGT |
| 36b4 | mouse | Fw: CCAGCGAGGCCACACTGCTG  Rv: ACACTGGCCACGTTGCGGAC |
| Cyclophilin | mouse | Fw: ATGGTCAACCCCACCGTGT  Rv: TTCTGCTGTCTTTGGAACTTTGTC |
| TNF | human | Fw: AGATGATCTGACTGCCTGGG  Rv: CTGCTGCACTTTGGAGTGAT |
| IL1β | human | Fw: AAGCCCTTGCTGTAGTGGTG  Rv: GAAGCTGATGGCCCTAAACA |
| IL10 | human | Fw: GCTGTCATCCATTTCTTCCC  Rv: CTCATGGCTTTGTAGATGCCT |
| 36B4 | human | Fw: TCATCAACGGGTACAAACGA  Rv: GCCTTGACCTTTTCAGCAAG |
| Cyclophilin | human | Fw: ACGGCGAGCCCTTGG  Rv: TTTCTGCTGTCTTTGGGACCT |
